# Extraembryonic endoderm cells induce neuroepithelial tissue in gastruloids

**DOI:** 10.1101/2020.02.13.947655

**Authors:** Noémie M. L. P. Bérenger-Currias, Maria Mircea, Esmée Adegeest, Patrick R. van den Berg, Marleen Feliksik, Mazène Hochane, Timon Idema, Sander J. Tans, Stefan Semrau

**Affiliations:** Leiden University, Einsteinweg 55, 2333 CC Leiden, Netherlands; Delft University of Technology, Department of Bionanoscience, Kavli Institute of Nanoscience, Van der Maasweg 9, 2629 HZ Delft, Netherlands; AMOLF, Science Park 104, 1098 XG Amsterdam, Netherlands

## Abstract

Stem-cell derived *in vitro* systems, such as organoids or embryoids, hold great potential for modeling *in vivo* biology and engineering living systems with novel functions. To unlock that potential, we need new ways to elicit higher-level organization. Here we show that adding extraembryonic endoderm (XEN) cells to mouse gastruloids leads to the formation of neural epithelia. By single-cell RNA-seq, imaging and differentiation experiments, we demonstrate the neural characteristics and spatial patterning of the epithelial tissue. We further show that the XEN cells differentiate reciprocally to a visceral endoderm-like state. Finally, we demonstrate that local inhibition of WNT signaling and production of a basement membrane by the XEN cells underlie the formation of the neuroepithelial tissue. In summary, we establish “XEN Enhanced Gastruloids” (XEGs) to explore heterotypic cellular interactions as a means to achieve complex, tissue-level organization *in vitro*.

## Introduction

Multicellular *in vitro* systems have become a major focus of biology and bioengineering over the last few years. Stem cell-derived systems, such as embryoids and organoids show complex organization and have the potential to exhibit functions that emerge from the interaction of multiple tissues^1–3^. These systems leverage differentiation mechanisms inherent to stem cells and rely heavily on self-organization. Their formation is therefore typically variable and difficult to control. By contrast, defined assemblies of differentiated cell types are much more controllable and can thus be engineered to exhibit novel functions^4^, but they are limited in complexity. To exploit the potential of stem cell-derived systems and eventually engineer them, we need to improve our control of these systems.

A promising starting point for complex, engineered systems are gastruloids. These aggregates of mouse or human embryonic stem cells (ESCs) recapitulate elements of embryonic development, such as embryo elongation and body axis formation^5–10^. Recently, tissue-level organization has been achieved by exogenous induction of relevant signaling pathways^11–13^. The possibility to induce various tissues in a controlled and reproducible way, would make gastruloids the ideal template for creating more complex systems.

Thus far, gastruloids are made exclusively from ESCs. *In vivo*, mammalian embryos receive important patterning inputs from extraembryonic cells, especially during the earliest embryonic stages^14^. Extraembryonic endoderm (XEN) cells have therefore been incorporated in embryoid systems^11, 15–17^. Remarkably, gastruloids achieve symmetry breaking without external cues from extraembryonic cells. This does not exclude, however, that XEN cells can unlock additional developmental potential and provide an opportunity to control gastruloid organization. XEN cells are derived from the primitive endoderm (PrE), which overlays the developing epiblast (see Fig. 1a for a schematic of early mouse development.) Subsequently, the PrE gives rise to the Parietal Endoderm (PE), which covers the inside of the blastocoel cavity^18^ and the Visceral Endoderm (VE), which surrounds the embryo until the formation of the visceral yolk sac and its integration in the embryonic gut^19, 20^. A subpopulation of the VE, the Anterior Visceral Endoderm (AVE) has a role in the establishment of the embryo’s body axes^21, 22^. Hence, XEN cells might have unexplored potential to pattern stem cell-derived model systems.

**Fig. 1.**
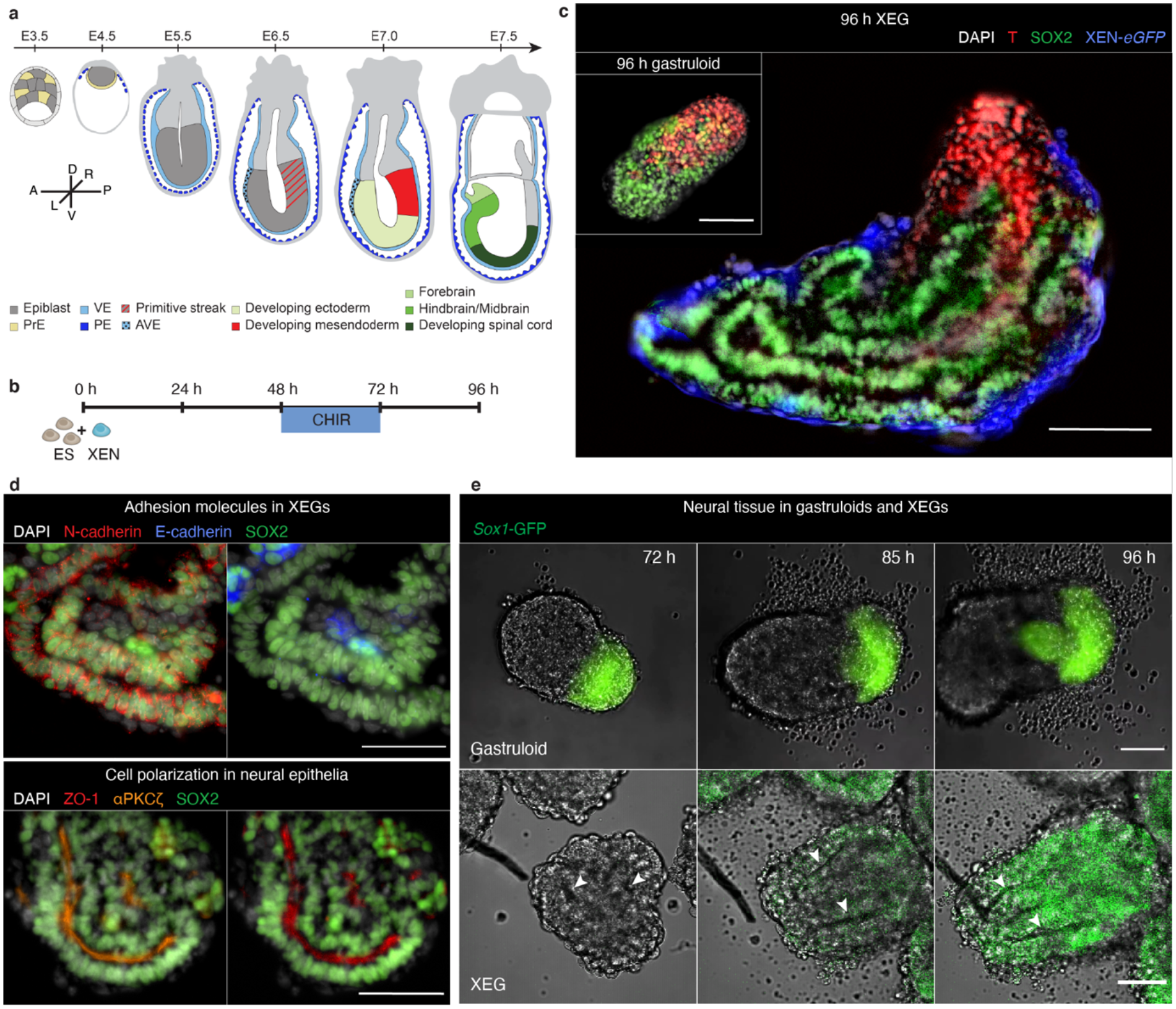
XEN cells induce neuroepithelial structures in XEN enhanced gastruloids (XEGs). **a**, Schematic of early mouse embryonic development. Tissues discussed in this manuscript are indicated with color. A: anterior, P: posterior, D: dorsal, V: ventral, L: left, R: right. **b**, Schematic of the culture protocol: at 0 h, 200 cells (150 ESCs and 50 XEN cells) were aggregated; CHIR was added between 48 h and 72 h after cell seeding; cell aggregates were cultured until 96 h. **c**, T and SOX2 expression in XEGs at 96 h (z-projection of whole mount immunostaining). Inset: expression in an aggregate resulting from the standard gastruloid protocol (without XEN cells) at 96 h. Scale bars: 100 µm. **d**, Expression of SOX2, E-cadherin, N-cadherin, ZO-1 and αPKCζ in XEGs at 96 h (immunostaining of cryosections). Scale bar: 50 µm. **e,** Live-cell imaging of SOX1 expression in a gastruloid (top panel, scale bar: 20 µm) and an XEG (lower panel, scale bar: 50 µm), grown with *Sox1-*GFP mESCs (see Supplementary Videos 3-8). In all images, a single z-plane is shown. The arrows indicate epithelial structures. **b, d,** Cell nuclei were stained with DAPI.

Here we report that aggregates of mouse embryonic stem cells (mESCs) and XEN cells can produce columnar neural epithelia. Using multiple markers, perturbation of the signaling pathways that play a role in neural development *in vivo*, and further differentiation to neural cell types, we confirmed that the epithelial structures indeed have neural characteristics. By single-cell RNA-seq, we identified differences in composition and molecular profiles between our new model system and regular gastruloids. We then established, that XEN-derived cells become visceral endoderm-like due to co-differentiation with the mESCs. Finally, we showed that XEN cells promote epithelia formation by local inhibition of primitive streak formation, likely via the WNT inhibitor DKK1, as well as through production of a basement membrane. Our study thus highlights the complex interplay between embryonic and extraembryonic cells and explores possible mechanisms underlying the great morphogenic potential of stem cell-derived systems

## Results

### XEN cells induce neuroepithelial structures in XEN enhanced gastruloids

We first implemented the original mouse gastruloid protocol^5^, in which mESCs are aggregated in N2B27 media and exposed to a pulse of WNT signaling for 24 h. After 96 h, this protocol results in elongated gastruloids. As reported before^5–7^, 96 h gastruloids contained localized primitive streak-like and neural progenitor-like compartments, marked by Brachyury (T) and SOX2, respectively (Fig. 1c, inset). Starting from the gastruloid protocol, we developed a new system by aggregating mESCs and XEN cells, keeping all other experimental conditions the same (Fig. 1b). We call our mixed aggregates “XEN Enhanced Gastruloids” (XEGs). Like gastruloids, 96 h XEGs showed an elongated morphology and localized T-positive and SOX2-positive compartments. However, unlike in gastruloids, SOX2+ cells in XEGs were organized in columnar epithelia surrounding one or several lumina (Fig 1c).

Expression of the broadly expressed neural marker SOX2 and the striking morphology suggested that the observed structures resemble neural epithelia. The lack of pluripotency marker expression (Supplementary Fig. 1a) excluded that the structures were formed by remaining undifferentiated cells. The presence of N-cadherin and absence of E-cadherin in the epithelia (Fig. 1d, top) is consistent with the known switch from E- to N-cadherin during neural differentiation *in vivo*^23^ and *in vitro*^24^. We could also observe that the epithelial cells were polarized and expressed apical markers ZO-1 and aPKC (Fig. 1d, bottom), consistent with neural epithelia *in vivo*^25^. Finally, we detected the neural progenitor markers PAX6 and NKX6.1^26^ in a subpopulation of epithelial cells (Supplementary Fig. 1b). Combined, these results suggest that the observed structures in XEGs have the characteristics of neural epithelia. To understand how these structures formed, we used time-lapse microscopy of developing XEGs. Around 48 h after seeding, cells formed rosette-like shapes (Supplementary Fig. 1c), which resembled structures found in Matrigel-embedded mESCs^27, 28^ and indicated a mesenchymal-epithelial transition. Subsequently, a columnar epithelium was formed. Then lumina opened at different places and merged between 48 h and 72 h (Supplementary Fig. 1d). During the final 24 h, the epithelium kept extending and differentiated further, as revealed by the expression of the neural progenitor marker SOX1^29^ (Fig. 1e, bottom). A SOX1 positive cell population also appeared in gastruloids within the same time frame, but, importantly, remained unorganized (Fig. 1e, top).

To explore the robustness of the protocol and identify optimal conditions for the formation of epithelial structures, we tested different ratios of mESCs and XEN cells (Supplementary Fig. 1e, f). Interestingly, even the smallest proportion of XEN cells tested (1:5), was able to induce some epithelia formation. On the other hand, elongation and symmetry breaking were inhibited when the proportion of XEN cells exceeded 1:2. A ratio of 1:3 gave optimal results, with the concurrence of SOX2-positive epithelia and T-positive cells in nearly all aggregates.

### Epithelia in XEGs show partial anteroposterior and dorsoventral patterning

To establish, how closely XEGs mimic *in vivo* development, we looked for patterning related to body axis formation. Like in gastruloids, expression of T (Fig. 1c), revealed a posterior-like compartment. Localized expression of anterior markers (ASCL1, PAX6, see Fig. 2a and Supplementary Fig. 1b) indicated an anterior-like compartment, although important canonical markers of the most anterior part of the embryo (OTX2, LEFTY1, EN1, ZIC1) could not be detected in XEGs (data not shown). The presence of anteroposterior patterning was supported by the expression of *Wnt4*, *Wnt8a* and *Fgf8* as gradients along the long axis of XEGs (Fig. 2b), which resembles the pattern found *in vivo*^30^. Neural epithelia in XEGs also expressed markers of both the dorsal (PAX6, MSX1) and ventral (NKX6.1, NRCAM) neural tube (Fig. 2c). While PAX6 and MSX1 were found in cells close to the outside of the XEG (which are usually in contact with the XEN cells), NKX6.1 and NRCAM were expressed broadly throughout the epithelial structures. In summary, 96 h XEGs show indications of anteroposterior and dorsoventral symmetry breaking.

**Fig. 2.**
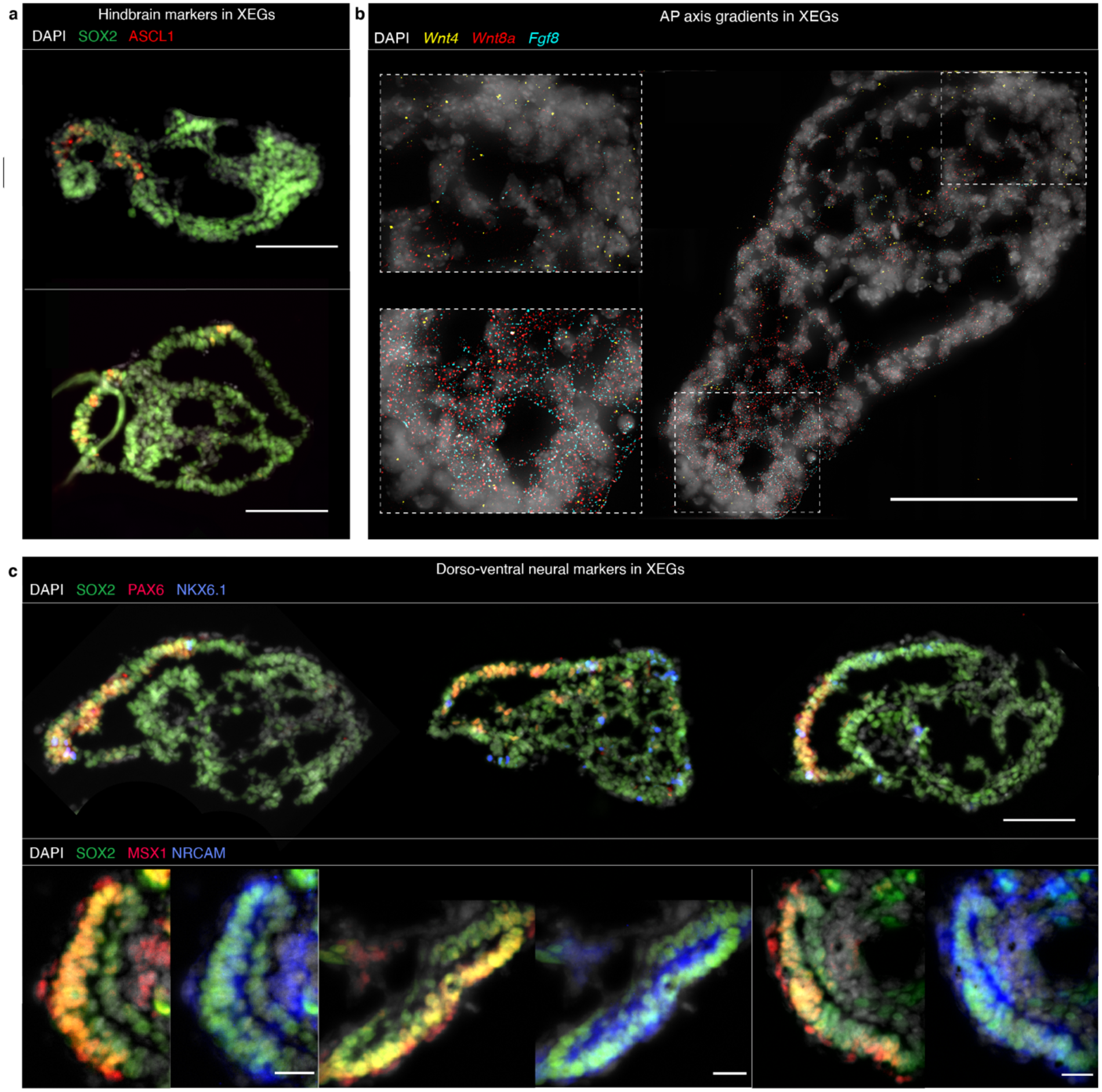
Neuroepithelia in XEGs show partial anteroposterior and dorsoventral patterning. **a**, Expression of SOX2 and ASLC1 in 96 h XEGs (immunostaining of sections). Scale bar: 100 µm. **b**, *Wnt4*, *Wnt8a* and *Fgf8* expression in XEGs at 96 h, visualized by smFISH on sections. Each diffraction limited dot is a single mRNA molecule. Scale bar: 100 µm. **c**, Expression of dorsal (PAX6, MSX1) and ventral (NKX6.1, NRCAM) neural tube markers in 96 h XEGs (immunostaining of sections). Top, expression of PAX6 and NKX6.1 Scale bar: 100 µm. Bottom, zoomed pictures of neural epithelia showing the expression of MSX1 and NRCAM. Scale bars: 20 µm. **a-c**, Cell nuclei were stained with DAPI.

### Signaling perturbation experiments and further differentiation support the neuroepithelial character

To further characterize the neuroepithelial structures, we tested how they respond to signaling inputs found *in vivo*. Specifically, we explored the response to BMP pathway inhibition, as well as Sonic Hedgehog (Shh) and retinoic acid (RA) pathway activation (Fig. 3a, b). The inhibition of BMP signaling, known to prevent premature neural specification^31^, resulted in an increased number of neural progenitor-like cells (marked by SOX2, PAX6 and NKX6.1). Sonic hedgehog, produced *in vivo* by the notochord and the floor plate (see schematic in Fig. 3a), is known to be necessary for the patterning of the ventral part of the neural tube^32^. The activation of the Hh signaling pathway led to a higher frequency of cells expressing a ventral marker (NKX6.1) in XEGs. RA, involved in anterior-posterior patterning^33^, strongly increased the number of cells expressing PAX6, which is found specifically in the anterior part of the neural tube^34^. The neuroepithelial structures in XEGs thus responded to signaling inputs as expected from *in vivo* development.

**Fig. 3.**
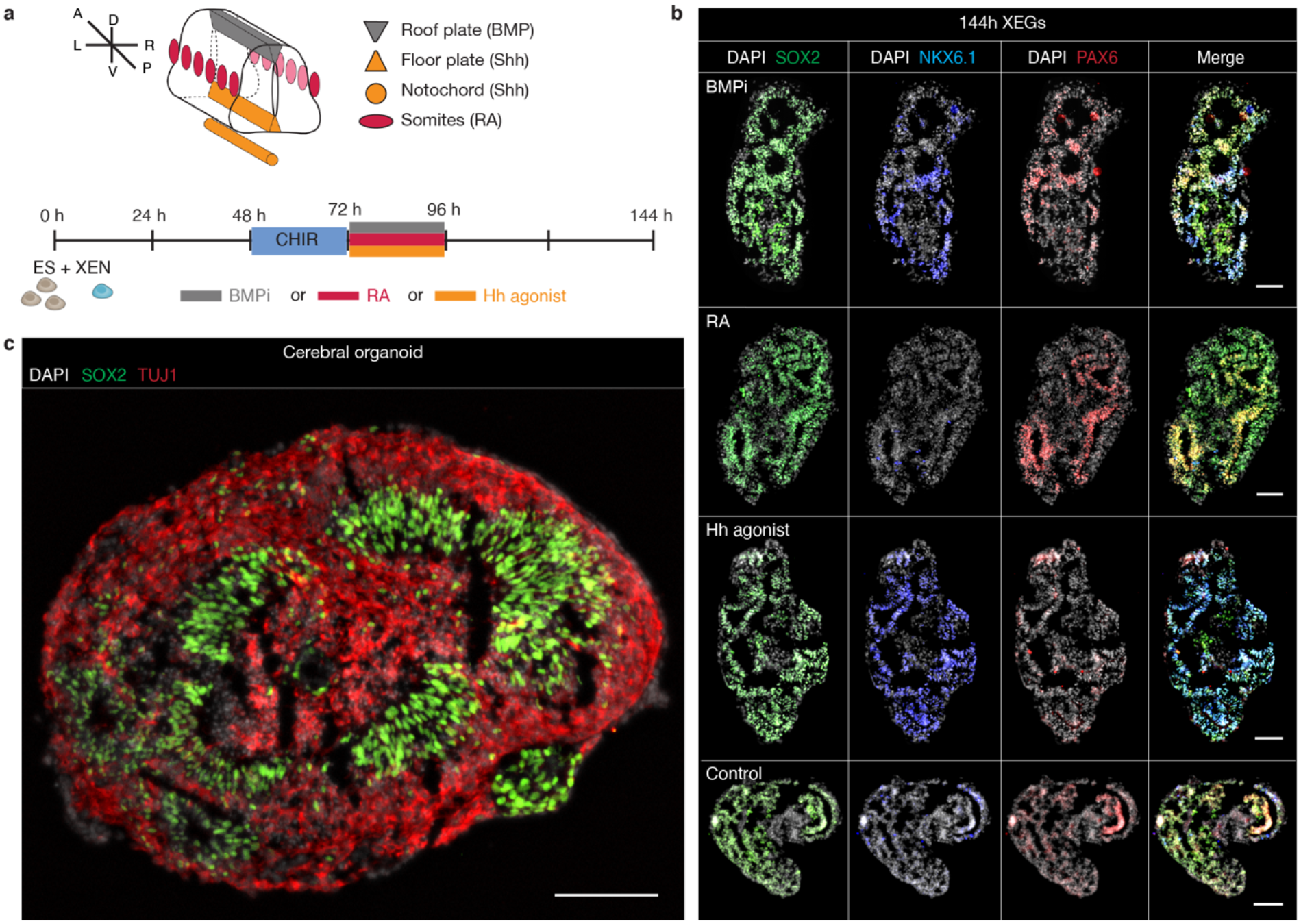
Signaling perturbation experiments and continued differentiation confirm neural character. **a,** Top: schematic of signaling sources patterning the developing neural tube *in vivo*. A: anterior, P: posterior, D: dorsal, V: ventral, L: left, R: right. Bottom: time line of the signaling experiments. XEGs were treated from 72 h to 96 h, with either BMP pathway inhibitor (BMPi), retinoic acid (RA) or hedgehog pathway agonist (Hh agonist). The XEGs were then allowed to grow for an additional 48 h before staining. **b,** Expression of SOX2, NKX6.1 and PAX6 in XEGs at 144 h, treated with the indicated factors (immunostaining of sections). n = 3 experiments. Scale bars: 100 µm. **c,** Expression of SOX2 and TUJ1 in XEGs, 8 days after cell seeding, differentiated according to a cerebral organoid protocol for 4 days (immunostaining of sections). Scale bar: 100 µm. **b-c**, Cell nuclei are stained with DAPI.

Even in the absence of RA, some hindbrain markers (ASCL1, PAX6) were sporadically expressed, typically concentrated at one pole of the XEG (Supplementary Fig. 1b, Fig. 2a). This hint of anterior specialization suggested a potential to differentiate further into cerebral tissue. Indeed, within four days of additional culture in cerebral organoid differentiation media^35^, XEGs developed a layered organization of neural progenitors (SOX2+/PAX6+) and neurons (TUJ1+/CTIP2+), surrounding cavities, reminiscent of tissue organization around brain ventricles (Fig. 3c, Supplementary Fig. 2a, b). Interestingly, we also observed a population of cells expressing the endothelial marker CD31 (Supplementary Fig. 2c). This might indicate that non-neural cells remained and might have differentiated further. Those CD31+ cells could specifically represent an early stage of vasculature.

### Single-cell RNA-seq reveals the transcriptional profiles of XEG cells

Having focused on the most striking, morphological difference between gastruloids and XEGs, we wanted to take a more comprehensive approach to reveal additional differences between the two model systems. To that end, we used single-cell RNA-sequencing (scRNA-seq) (Supplementary Fig. 3a-e). By mapping the data to single-cell transcriptomes of mouse embryos from E6.5 to E8.5^36^ (Supplementary Fig. 4a, b) we classified the transcriptional identity of the cells (Fig. 4a, b). Except for the least abundant cell types, the distribution of cell types was consistent across two biological replicates (Fig. 4c). Expression of known markers confirmed the classification by mapping to *in vivo* data (Supplementary Fig. 4c, Supplementary Table 1). Most cell types belonged to the E8.0 or E8.5 embryo (Supplementary Fig. 4d), which might indicate that *in vitro* differentiation proceeded roughly with the same speed as *in vivo* development.

**Fig. 4.**
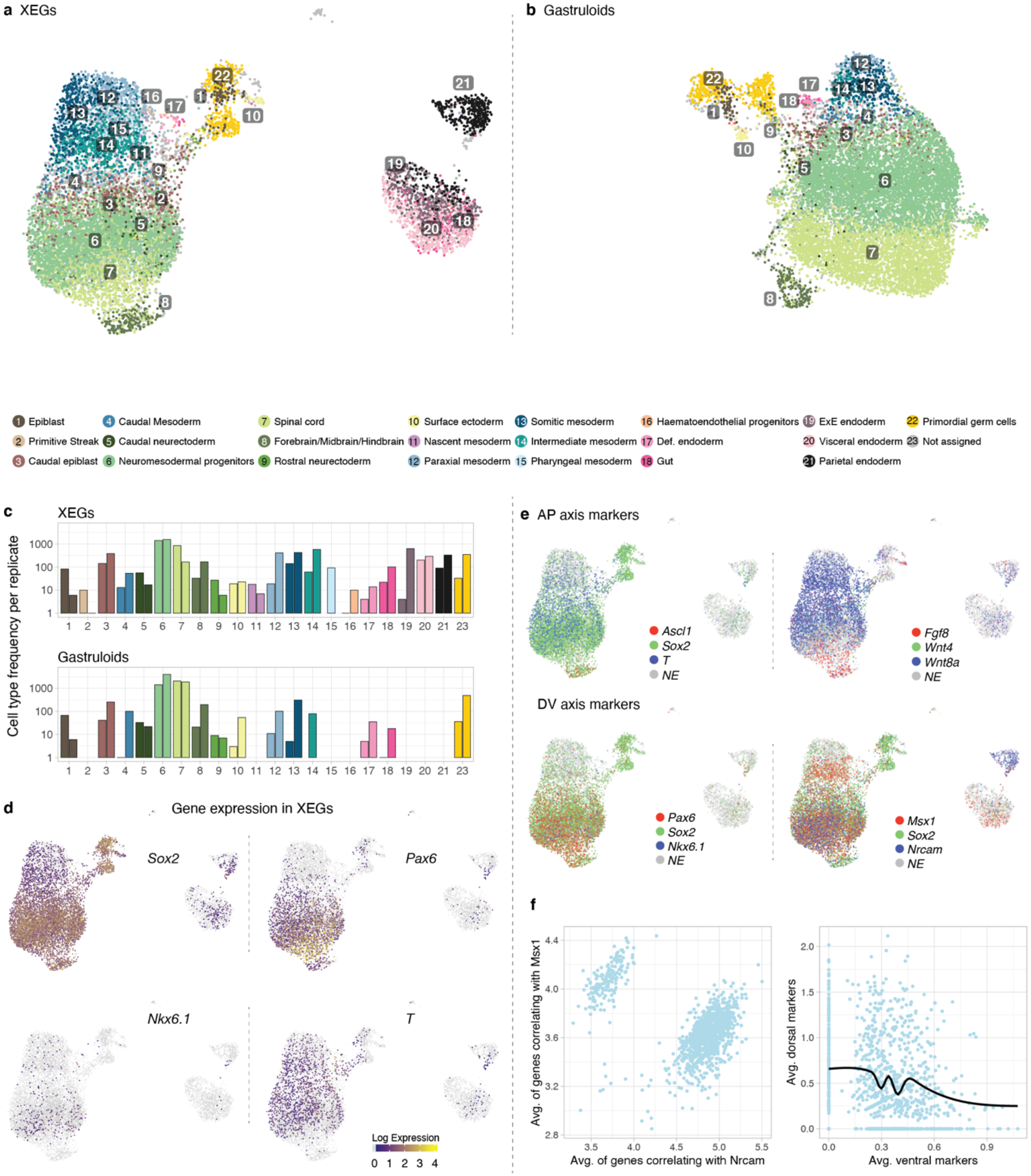
Single-cell RNA-seq reveals the transcriptional profiles of XEG cells. **a,b**, UMAP of cells in XEGs and gastruloids colored by cell type based on the mapping to *in vivo* data shown in Supplementary Fig. 4a,b. **c**, Cell type frequencies for both replicates in XEGs and gastruloids. **d**, *Sox2*, *Pax6*, *Nkx6.1* and *T* log-expression levels indicated by color in UMAPs of XEGs. **e**, Top, UMAP of cells in XEGs with log expression of genes expressed along the anteroposterior (AP) axis indicated by colors. Left: *Aslc1*, *Sox2* and *T*, right: *Wnt4*, *Wnt8a* and *Fgf8*. Bottom, UMAP of cells in XEGs with log expression of genes expressed along the dorsoventral (DV) axis indicated by colors. Left: *Pax6*, *Sox2* and *Nkx6.1*, right: *Msx1*, *Sox2* and *Nrcam* indicated by color. All UMAPs show both replicates, batch corrected. **f**, Left, scatter plot of the average expression of the top 10 genes correlating most strongly with *Msx1* and *Nrcam*, respectively, in our scRNA-seq data. Right, scatter plot of the average expression of 10 canonical marker genes for dorsal and ventral neural progenitors, respectively, with the moving average indicated by the black solid line (see Methods for the gene lists).

Neuromesodermal progenitors (NMPs) and spinal cord-like cells were the most abundant in both model systems (Fig. 4c). Gastruloids thus already contain cells of the neural lineage, which, however, seem to lack organization (Fig. 1c, inset). To identify the cells forming epithelial structures in XEGs, we used the neural markers *Sox2*, *Pax6* and *Nkx6.1*^26^, which we had detected by immunostaining (Supplementary Fig. 1b). We found these markers to be co-expressed in cells classified as “spinal cord” and “brain” in the scRNA-seq data (Fig. 4d, Supplementary Fig. 4e), confirming their neural ectoderm identity. While NMPs also expressed *Sox2* and *Nkx6.1*, neuroepithelial structures where clearly distinguishable by the presence of *Pax6* and the absence of *T*.

The scRNA-seq results also supported the existence of anteroposterior, as well as dorsoventral patterning. Several anterior and posterior markers were clearly expressed in separate subpopulations (Fig. 4e). Differential gene expression analysis between “spinal cord” cells and other cells in XEGs identified markers of both the dorsal and ventral neural tube (Fig. 4e, Supplementary Fig. 5a, Supplementary Table 2). Two sets of genes that were either co-expressed with *Msx1* or *Nrcam*, respectively, showed strong anticorrelation (Fig. 4f, left), which confirmed our immunostaining results (Fig. 2c). Similarly, the averages of canonical dorsal and ventral markers were anti-correlated (Fig. 4f, right), despite individual markers being lowly expressed. A comparison between XEGs and gastruloids revealed that neural ectoderm-like cells expressed more dorsal markers in XEGs (Supplementary Fig. 5b). This dorsal identity was confirmed by mapping the neural ectoderm-like cells to single-cell expression profiles of *in vivo* neural tube^37^ (Supplementary Fig. 5c). The majority of neural ectoderm-like cells from XEGs turned out to be more similar to dorsal progenitors *in vivo*. All in all, the scRNA-seq results support dorsoventral patterning in XEGs, with a bias towards the dorsal identity, compared to gastruloids.

Overall, both model systems showed a diverse cell type distribution, comprising a variety of mesodermal as well as anterior cell types. Thus, XEG cells are not globally biased towards the neural fate, as occurring in other protocols for induction of neural epithelia^12, 28, 38^. On the contrary, XEGs even contained a bigger proportion of mesoderm-like cells. While paraxial, intermediate and somitic mesoderm were present in both model systems, only XEGs contained primitive streak, nascent mesoderm, pharyngeal mesoderm and hematoendothelial progenitors (Fig. 4c, Supplementary Fig. 6a). To confirm the presence of mesodermal cell types in XEGs, we focused on two genes, *Tbx6* and *Pax2*, which are markers of nascent and intermediate mesoderm, respectively. Our single-cell RNA-seq data showed expression of both genes in subpopulations of XEG cells (Supplementary Fig. 6b) and immunostaining confirmed their presence (Supplementary Fig. 6c). However, we did not observe any tissue-level organization of those cells in XEGs. Taken together, these results suggest that the XEN-derived cells in XEGs also have an effect on the mesodermal cell population, which, in turn, could potentially pattern the neuroepithelial tissues.

### Most XEN cells become VE-like in XEGs

Compared to gastruloids, XEGs additionally contained extraembryonic endoderm cell types (Fig. 4c). By using GFP-expressing XEN cells in XEGs (Supplementary Fig 7a), we established that those cell types were exclusively differentiated from XEN cells. By comparison to undifferentiated XEN cells, which were spiked into the scRNA-seq samples, we studied the transcriptional changes in XEN-derived cells. Undifferentiated XEN cells mostly mapped to PE^36^ (Fig. 5a, Supplementary Fig. 7b), consistent with a previous study^39, 40^. Their derivatives in XEGs mapped to multiple kinds of extraembryonic endoderm: PE, embryonic VE and extraembryonic VE. Interestingly, some also mapped to gut, reminiscent of the contribution of VE to the gut *in vivo*^19, 41^. The identification of those cell types was confirmed by mapping our scRNA-seq data to an endoderm-focused scRNA-seq dataset^41^ (Supplementary Fig. 7c,d). Quantification revealed that, on average, 8% of the initially PE-like XEN cells acquired a gut-like and 66% a VE-like transcriptomic profile (29% embryonic VE, 37% extraembryonic VE) when co-cultured in XEGs. However, 25% retained a PE-like transcriptome (Fig. 5a). Differential gene expression analysis between undifferentiated XEN cells and XEN-derivatives revealed several differences (Supplementary Fig. 7e, Supplementary Table 3). PE markers were less expressed in XEN-derivatives, while VE markers were highly expressed, suggesting that most XEN cells differentiate from at PE to a VE-like state in XEGs. To validate this finding experimentally, we performed single-molecule FISH of the PE marker *Fst*^42^, the VE marker *Spink1*^43^ and the pan-extraembryonic endoderm marker *Dab2*^44^ (Fig. 5b). This measurement showed that XEN-derived cells in XEGs only expressed *Dab2* and *Spink1*, while undifferentiated XEN cells broadly co-expressed all markers. Some XEN cells in XEGs were also highly expressing E-cadherin, known to be expressed in VE^45^ (Fig. 5c). However, the more anterior VE marker *Hhex*^46^ was not detected by single-molecule FISH (Fig. 5d). Exposing undifferentiated XEN cells to CHIR in the same way as XEGs did not cause differentiation (Fig. 5d), which suggests that the interaction with mESCs plays a role.

**Fig. 5.**
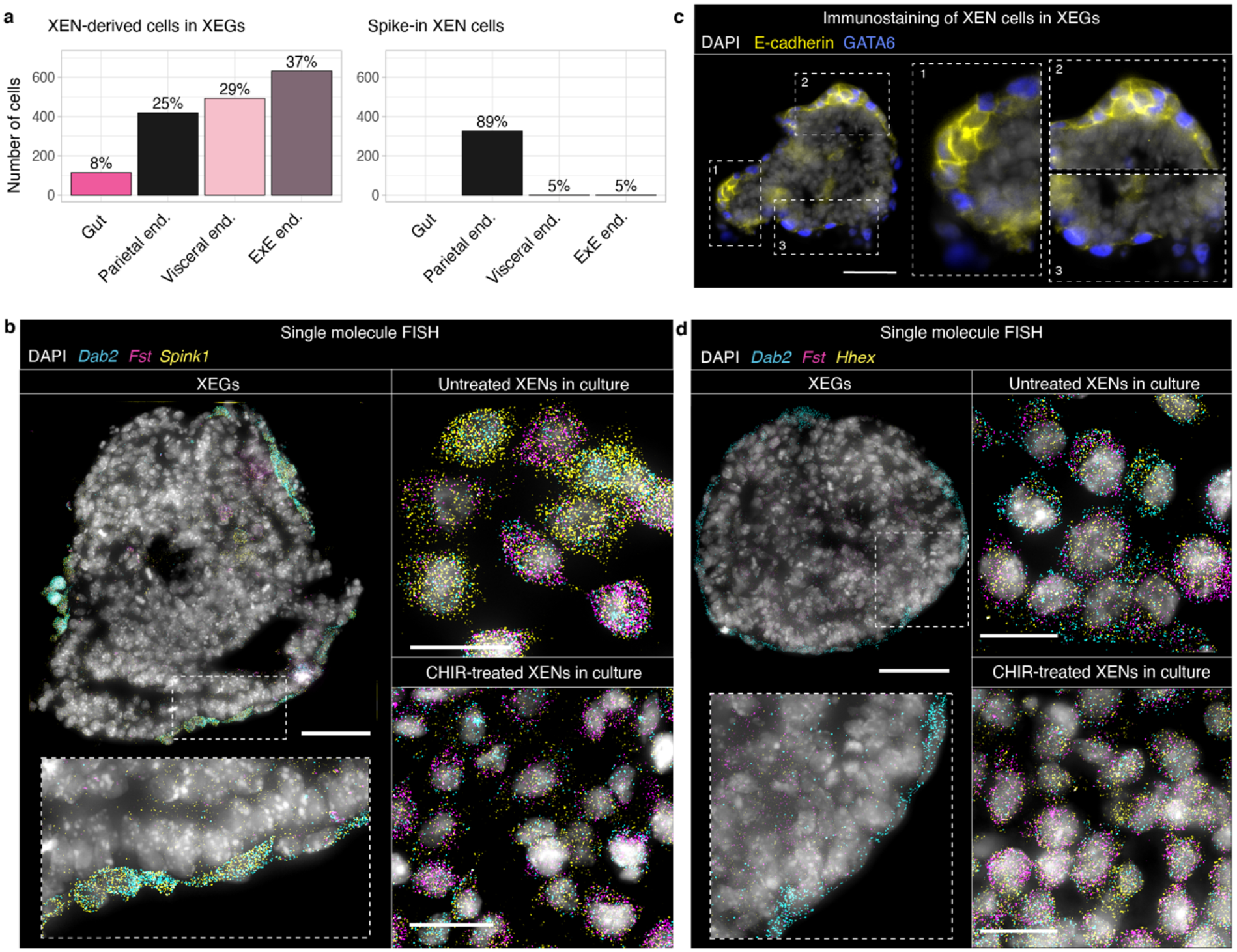
Most XEN cells become VE-like in XEGs. **a**, Left, cell types of XEN-derived cells in XEGs. Cells were classified as gut, PE (“parietal end.”), embryonic VE (“visceral end.”) or extraembryonic VE (“ExE end.”) by mapping to the data set from the Pijuan-Sala et al. dataset^36^. Right, cell types of spiked-in XEN cells. **b**, *Dab2*, *Spink1* and *Fst* expression in a section of an XEG at 96 h (left, scale bar: 50 µm), in XEN cells cultured under standard maintenance conditions (top right, scale bar: 20 µm) and in XEN cells treated with CHIR according to the XEG protocol (bottom right, scale bar: 20 µm). Expression was visualized by smFISH. Each diffraction limited dot is a single mRNA molecule. **c**, Expression of E-cadherin in XEGs at 96 h (immunostaining of sections). XEN cells were localized by expression of GATA6. Scale bars: 50 µm. **d**, *Dab2*, *Fst* and *Hhex* expression in a section of an XEG at 96 h (left, scale bar: 50 µm), in XEN cells cultured under standard maintenance conditions (top right, scale bar: 20 µm) and in XEN cells treated with CHIR according to the XEG protocol (bottom right, scale bar: 20 µm). Expression was visualized by smFISH. Each diffraction limited dot is a single mRNA molecule. **b-d**, Cell nuclei were stained with DAPI. The dashed boxes are shown at a higher magnification in the insets.

Taken together, these results suggest that XEGs mimic elements of embryo organization *in vivo*, where the VE is in direct contact with the embryo proper. While undifferentiated XEN cells have both PE and VE characteristics, the majority (66%) of these cells becomes more VE-like, due to the presence of mESCs.

### XEN cells guide symmetry breaking by local inhibition of WNT signaling

Although XEN-derived cells in XEGs did not express canonical AVE markers (Supplementary Fig. 4c), we were wondering if they might effectively carry out an AVE-like patterning function. XEN cells always formed the outermost layer (Fig. 1b, Supplementary Fig. 1f), resembling *in vivo* organization. Focusing on XEGs partially covered with XEN cells, we observed that epithelial structures were always adjacent to the XEN cells, while the T-positive population was on the opposite side (Supplementary Fig. 1f). Notably, this organization was already established at 72 h, when aggregates are still spherical (Fig. 6a). This observation suggested that XEN cells guide the symmetry breaking by a local effect on the adjacent mESCs. We speculated that this effect might be mediated by a basement membrane produced by the XEN cells. As established above, cells were polarized early during XEG development, prior to forming a columnar epithelium (Supplementary Fig. 1c, d). This epithelium was supported by a basement membrane containing laminin and collagen (Fig. 6b, c) which were mostly produced by the XEN cells (Supplementary Fig. 8a, top-right). It has been demonstrated previously, for small aggregates of mESCs, that the presence of an extracellular matrix can be sufficient for polarization and lumen formation^27, 28, 38^. Growing gastruloids in Geltrex did induce the formation of inner cavities; however, we did not observe a columnar epithelium (Supplementary Fig. 8b).

**Fig. 6.**
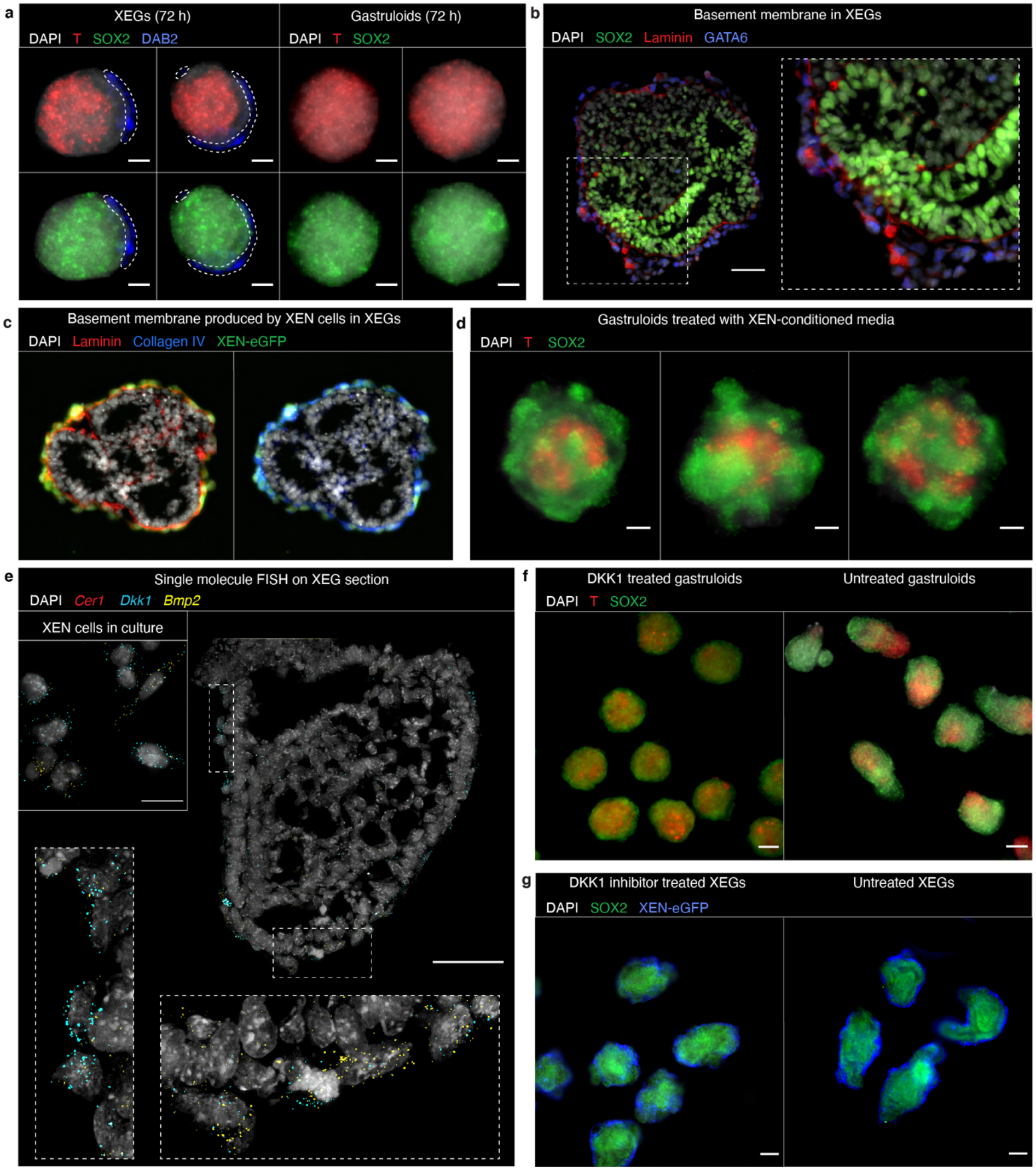
XEN cells guide symmetry breaking by locally inhibiting primitive streak formation. **a**, T and SOX2 expression in XEGs (left) and gastruloids (right) at 72 h (z-projection of whole mount immunostaining). XEN cells were localized by expression of DAB2 and are indicated by a dashed outline. **b**, Expression of SOX2 and laminin in XEGs at 96 h (immunostaining of sections). XEN cells were localized by expression of GATA6. The dashed boxes are shown at a higher magnification in the insets. Scale bars: 50 µm. **c**, Expression of collagen IV and laminin in XEGs at 96 h (immunostaining of sections). XEN cells were localized by expression of H2B-GFP. Scale bars: 100 µm. **d**, T and SOX2 expression in gastruloids grown in XEN-conditioned media at 96 h (z-projection of whole mount immunostaining). **e**, *Cer1*, *Dkk1* and *Bmp2* expression in a section of a XEG at 96 h (scale bar: 50 µm) or XEN cells cultured under standard maintenance conditions (inset, scale bars: 20 µm). Expression was visualized by smFISH. Each diffraction limited dot is a single mRNA molecule. **f**, T and SOX2 expression in 96 h gastruloids treated with 200 ng/mL DKK1 and untreated (z-projection of whole mount immunostaining). Scale bars: 100 µm. **g**, T and SOX2 expression in 96 h XEGs treated with 0.25 µM DKK1 inhibitor WAY-262611 and untreated (z-projection of whole mount immunostaining). Scale bars: 100 µm. **a-g**, Cell nuclei were stained with DAPI.

Since an extracellular matrix was not sufficient to obtain neuroepithelial structures, we were wondering, which other mechanisms might play a role. We observed that gastruloids grown in medium conditioned by XEN cells, did not elongate and had a T-positive cell population that was restricted to the center of the aggregate (Fig. 6d). Thus, we reasoned that diffusible factors produced by the XEN cells might have a role in the formation of the neuroepithelial structures. We hypothesized that the WNT inhibitor DKK1, which is expressed by the XEN cells (Fig. 6e; Supplementary Fig. 8a, bottom-right), might be one of those factors. *In vivo*, DKK1 is expressed by the AVE and limits the growth of the primitive streak^47^. Notably, other factors known to act in this process (CER1, LEFTY1) are not expressed in XEGs (Fig. 6e, Supplementary Fig. 4c). Growing gastruloids in the presence of DKK1 resulted in a round morphology, with the T-positive cells confined to the center, as observed for XEN-conditioned medium (Fig. 6f, Supplementary Fig. 8d). Factors limiting the primitive streak expansion *in vivo* are also known to preserve the anterior part of the epiblast and are thereby necessary for proper ectoderm domain differentiation^48^. Thus, we wanted to explore, if DKK1 could have a similar role in XEGs and bias differentiation towards the ectodermal lineage. Growing XEGs with the DKK1 inhibitor WAY-262611^49^ led to a suppression of epithelial structures (Fig. 6g, Supplementary Fig. 8e). Notably, XEGs treated with the DKK1 inhibitor resembled gastruloids, showing elongated shapes but no epithelial structures. Since growing XEGs without CHIR resulted in similar epithelial structures as in regular XEGs (Supplementary Fig. 8c), XEN cells likely suppress pre-existing, low-level endogenous WNT activity^6, 50^. Finally, we noticed that XEN-derived cells highly express BMPs (Supplementary Fig. 8a, bottom-right) and that several of the dorsal markers expressed in XEGs, are induced by BMP signaling (Supplementary Table 4). Thus, XEN-derived cells might also contribute to the dorsal patterning of the neural progenitors-like cells in XEGs.

All combined, our experiments suggest that XEN cells guide symmetry-breaking by local inhibition of cell differentiation into a T-positive population. Diffusible factors, including DKK1, and the presence of a basement membrane both seem to contribute to the formation of the neuroepithelial structures.

## Discussion

In this study we explored how the interaction between embryonic and extraembryonic cells in a multicellular *in vitro* system can lead to the formation of neuroepithelial tissue. The observed dynamics and organization of the neuroepithelia in XEGs is clearly different from the formation of the neural tube *in vivo*, which occurs via two distinct mechanisms^51^. During primary neurulation, the main part of the neural tube is formed by the folding of the neural plate, an epithelial sheet of neural ectoderm cells. Secondary neurulation, which gives rise to the most posterior part of the neural tube, works differently: mesenchymal cells condense to an epithelial rod which cavitates to form a tube^52, 53^. The two parts of the tube are then connected during junctional neurulation^54^. While we did not observe cell rearrangements characteristic of primary neurulation, the rosette formation seen in XEGs was reminiscent of secondary neurulation^51^, which gives rise to the posterior neural tube. Nevertheless, we were able to differentiate XEGs towards neural tissue that showed layered organization reminiscent of the cortex, which derives from the anterior neural tube. The neuroepithelia in 96 h XEGs are therefore likely still plastic and would need additional time and signaling inputs to become fully specified. The recently reported Trunk-Like Structures (TLS)^55^, another gastruloid-derived model system, also produce neural tube-like tissues, together with mesodermal tissue resembling somites. Notably, TLS are formed exclusively from mESCs and are grown in Matrigel with additional exogenous signaling cues (BMP signaling inhibitor LDN193189 and CHIR). Another recent approach uses BMP4-treated ESCs as signaling centers to elicit neural tube-like tissues and other embryonic structures in untreated ESCs^13^.

The fact that the majority of XEN cells becomes VE-like in XEGs clearly shows that there are reciprocal interactions between the co-differentiating, embryonic and extraembryonic cells. This observation supports the notion that such interactions are necessary for proper development, as previously observed *in vivo*^19, 56^. Thus, XEN cells represent a potential way to augment existing developmental *in vitro* systems, by providing a basement membrane and extraembryonic signaling inputs.

Finally, with their large diversity of cell types, XEGs could be a starting point for developing more complex models containing all three germ layers as well as extraembryonic cells. Specifically, the CD31 positive endothelial cells observed in the cerebral organoids obtained from XEGs might be able to form a vascular network if additional signaling cues are given^57^.

In conclusion, in this study we showed how the morphogenic potential of heterotypic cell-cell interactions can be leveraged to enhance the organization of multicellular *in vitro* systems.

## Methods

### Experimental methods

#### Cell culture

All cell lines were routinely cultured in KO DMEM medium (Gibco) supplemented with 10% ES certified FBS (Gibco), 0.1 mM 2-Mercaptoethanol (Sigma-Aldrich), 1 × 100 U/mL penicillin/streptomycin, 1x MEM Non-Essential Amino Acids (Gibco), 2 mM L-glutamine (Gibco), 1000 U/mL mouse LIF (ESGRO). Cells were passaged every other day and replated in tissue-culture treated dishes coated with gelatin. E14 mouse ES cells were provided by Alexander van Oudenaarden. The *Sox1^GFPiresPac^* mouse ES cell line was created by Mario Stavridis and Meng Li in the group of Austin Smith^58^ and provided by Sally Lowell. XEN and XEN-*eGFP* were provided by Christian Schröter^40^. All cell lines were regularly tested for mycoplasma infection. The ES-mCherry-GPI cell line was obtained by introducing a mCherry-GPI transgene in the *Pdgfra^H2B-GFP^* cell line, provided by the group of Anna-Katerina Hadjantonakis^59^.

### Differentiation

#### Gastruloids

The gastruloid differentiation protocol was adapted from van den Brink et al. ^5^. ES cells were collected from tissue-culture treated dishes by trypsinization, gentle trituration with a pipet and centrifugation (1200 r.p.m., 3 min). After collection, cells were resuspended in 2 mL of freshly prepared, prewarmed N2B27 medium: DMEM/F12 (Life technologies) supplemented with 0.5 × N2 supplement (Gibco), 0.5 × B27 supplement (Gibco), 0.5 mM L-glutamine (Gibco), 1 × 100 U/mL penicillin/streptomycin (Gibco), 0.5 × MEM Non-Essential Amino Acids (Gibco), 0.1 mM 2-Mercaptoethanol (Sigma-Aldrich). Cells were counted to determine the cell concentration. For gastruloids, 200 ES cells were seeded in 40 µL of N2B27 in each well of a round-bottom low-adherence 96-well plate. 48 h after seeding, 150 µL of prewarmed N2B27 supplemented with 3 µM of GSK3 inhibitor (CHIR99021, Axon Medchem) was added to each well. 72 h after seeding, 150 µL of medium was removed from each well and replaced by 150 µL of preheated N2B27. Gastruloids were collected at 96 h after seeding and fixed with 4% paraformaldehyde (PFA, Alfa Aesar) overnight at 4 °C.

For the experiments with gastruloids grown in Geltrex, cell aggregates were collected at 24 h, 48 h and 72 h and embedded into LDEV-Free, hESC-Qualified, reduced growth factor Geltrex (Gibco) in culture dishes for the rest of the procedure. Only the gastruloids transferred at 72 h showed robust growth. At 96 h, culture dishes were covered with ice-cold PBS and placed on a shaker at 4 °C for 10 min. Gastruloids were gently collected by pipetting and washed three times by centrifugation in ice-cold PBS to remove the gel, then fixed with 4% PFA overnight at 4 °C.

#### XEN Enhanced Gastruloids (XEGs)

ES and XEN cells were collected from tissue-culture treated dishes by trypsinization, gentle trituration with a pipet and centrifugation (1200 r.p.m., 3 min). After collection, cells were resuspended in 2 mL of fresh and prewarmed N2B27 medium. Cells were counted to determine cell concentration. For XEGs, several ratios of XEN and ES cells were tested (1:1, 1:2, 1:3, 1:4, 1:5) and compared with the regular gastruloid condition (0:1). The total number of cells was fixed at 200. Over two separate experiments, the proportion of organoids showing T staining and epithelial structures was quantified (total number of embryonic organoids 1:1=179, 1:2=143, 1:3=143, 1:4=140, 0:1=130) and the optimal ratio was determined to be 1:3 (Supplementary Fig. 1e,f). A total of 200 cells (150 ES cells and 50 XEN cells) was seeded in 40 µL of N2B27 in each well of a round-bottom low-adherence 96-well plate. 48 h after seeding, 150 µL of prewarmed N2B27 supplemented with 3 µM of GSK3 inhibitor (CHIR99021, Axon Medchem) was added to each well. 72 h after seeding, 150 µL of medium was removed from each well and replaced by 150 µL of prewarmed N2B27. XEGs were collected at 96 h after seeding and fixed with 4% PFA overnight at 4 °C. For the experiment of XEGs grown without GSK3 inhibitor, cells were seeded as usual. At 48 h, 150 µL of preheated N2B27 was added to each well. At 72 h, 150 µL of medium was removed from each well and replaced by 150 µL of prewarmed N2B27. XEGs were collected at 96 h after seeding.

For the smFISH control experiments, XEN cells were seeded at low density in N2B27 medium. At 48 h the medium was replaced by prewarmed N2B27 supplemented with 3 µM of GSK3 inhibitor. 72 h after seeding, the medium was replaced with prewarmed N2B27. Cells were fixed at 96 h with 4% PFA for 1 h at 4 °C.

#### Neural differentiation

For neural differentiation, a protocol for creating cerebral organoids was adapted from Lancaster et al. ^35^. Instead of collecting XEGs at 96 h, the medium was replaced by cerebral organoid differentiation medium: DMEM-F12 (Life technologies), Neurobasal (Gibco), 0.5 × B27 supplement containing vitamin A (Gibco), 0.5 × N2 supplement (Gibco), 2.5 µM/mL Insulin, 2mM L-glutamine (Gibco), 0.5 × MEM-Non-Essential Amino Acids (Gibco), 1 × 100 U/mL penicillin-streptomycin and 0.05 mM 2-Mercaptoethanol (Sigma-Aldrich). At 168 h, aggregates were collected and transferred, with fresh medium, into 10 cm dishes on an orbital shaker installed in the incubator (85 r.p.m.). Aggregates were grown until 192 h (8 days) during which medium was refreshed every other day until collection. Collected aggregates were fixed with 4% PFA for 48 h at 4 °C.

#### Signaling experiments

In the signaling experiments with XEGs, aggregates were treated between 72 h and 96 h with either LDN193189 (BMPi, 100 nM, Reagents Direct), a potent BMP pathway inhibitor, Purmorphamine (1 µM, STEMCELL Technologies), a small molecule agonist of the hedgehog pathway, Retinoic acid (RA, 100 nM, Sigma-Aldrich) or DMSO (0.1% final concentration, Sigma Aldrich) as a vehicle control. For this experiment, the XEGs were allowed to grow for an additional 48 h before fixation (144 h total growth) and preparation for staining (see Immunostaining).

DKK1 signaling pathways perturbation was performed in two ways, using DKK1 (Sigma-Aldrich) for activation in gastruloids and Way262611 (Sigma-Aldrich) for inhibition in XEGs. Gastruloids and XEGs were seeded according to the usual protocols. At 24 h, 40 µL of N2B27 supplemented with various concentration of DKK1 or Way262611 respectively, were added to each well. Next steps of the protocol were performed using N2B27 supplemented with DKK1 or Way262611. Aggregates were fixed at 96 h with 4% PFA overnight at 4 °C.

### Immunostaining

#### Fixation and blocking

After collection, gastruloids and XEGs were fixed in 4% PFA at 4 °C overnight. Tissue resulting from the cerebral organoid protocol was fixed under the same conditions, but for 48 h. After fixation, samples were washed three times in washing solution (PBS, 1% bovine serum albumin (BSA)) and incubated at 4 °C in blocking buffer (PBS, 1% BSA, 0.3% Triton-X-100) for a minimum of 16 h. Samples for smFISH were washed 3 times in PBS after fixation and stored in 70% ethanol at 4 °C. To stain E14 cells for pluripotency markers, cells in suspension were fixed for 30 min in 4% PFA at 4 °C, washed three times in washing solution at RT and incubated in blocking buffer for 1 h at 4 °C.

#### Whole-mount immunolabeling and clearing

Immunolabeling and clearing of gastruloids and XEGs were based on the protocol described by Dekkers et al., ^60^. Briefly, after fixation and blocking, samples were incubated with primary antibodies at 4 °C overnight on a rolling mixer (30 r.p.m.) in organoid washing buffer (OWB) (PBS, 2% BSA, 0.1% Triton-X-100) supplemented with 0.02% sodium dodecyl sulfate (SDS), referred to as OWB-SDS. The following primary antibodies were used: rat anti-SOX2 (1:200, 14-9811-82, Thermo Fisher Scientific), goat anti-T (1:200, sc-17745, Santa Cruz Biotechnology), goat anti-T (1:100, AF2085, R&D systems), mouse anti-DAB2 (1:100, 610464, BD Biosciences). The next day, samples were washed three times for 2 h in OWB-SDS at RT, followed by incubation with secondary antibodies (donkey anti-goat Alexa Fluor 488 (1:200, A-11055, Thermo Fisher Scientific), donkey anti-rat Alexa Fluor 488 (1:200, A-21208, Thermo Fisher Scientific), donkey anti-goat Alexa Fluor 555 (1:200, A-21432, Thermo Fisher), donkey anti-mouse Alexa Fluor 555 (1:200, A-31570, Thermo Fisher Scientific), chicken anti-rat Alexa Fluor 647 (1:200, A-21472, Thermo Fisher Scientific)) and 4’,6-diamidino-2-phenylindole (DAPI, 1 µg/mL, Merck) in OWB-SDS at 4 °C overnight on a rolling mixer (30 r.p.m.), protected from light. Finally, samples were washed three times for 2 h in OWB-SDS at RT. Clearing was performed by incubation in fructose-glycerol clearing solution (60% vol/vol glycerol, 2.5 M fructose) for 20 min at RT. Samples were imaged directly after clearing or stored at 4 °C in the dark.

#### Cryosectioning and immunolabeling of sections

Prior to cryosectioning, fixed and blocked samples were incubated sequentially in sucrose solutions (10, 20 and 30%) for 30 min (gastruloids and XEGs) or 2 h (cerebral organoids) at 27 °C, and embedded in optimal cutting temperature (OCT) compound. Samples in OCT were placed on dry ice for rapid freezing, and stored at −80 °C prior to cryosectioning. Samples were cut to cryosections (10 µm thickness) using a cryostat (Thermo Fisher Scientific, USA) and cryosections were placed on poly-L-lysine coated glass slides (Merck). The slides were stored directly at −80 °C. For immunofluorescence staining, slides were thawed and rinsed with PBS for 10 min at RT to dissolve the OCT. Subsequently, slides were incubated overnight at 4 °C with the following primary antibodies diluted in blocking buffer: rat anti-SOX2 (1:200, 14-9811-82, Thermo Fisher Scientific), goat anti-T (1:200, sc-17745, Santa Cruz Biotechnology), mouse anti-N-cadherin (1:200, 33-3900, Thermo Fisher Scientific), rabbit anti-E-cadherin (1:200, 3195, Cell Signaling Technology), rabbit anti-PAX6 (1:100 (cerebral organoids) or 1:200 (gastruloids, XEGs), 42-6600, Thermo Fisher Scientific), mouse anti-NKX6.1 (1:200, F55A12, Developmental Studies Hybridoma Bank), rabbit anti-NKX6.1 (1:200, HPA036774, Merck), mouse anti-TUJ1 (1:200, 801202, BioLegend), rabbit anti-CD31 (1:50, ab28364, Abcam), rabbit anti-GATA6 (1:200, PA1-104, Thermo Fisher Scientific), goat anti-GATA6 (1:200, AF1700, R&D Systems), rabbit anti-Laminin (1:200, PA1-16730, Thermo Fisher Scientific), mouse anti-OCT4 (1:200, MA1-104, Thermo Fisher Scientific). The next day, the slides were washed twice for 10 min in PBS at RT. Subsequently, the slides were incubated with secondary antibodies (donkey anti-goat Alexa Fluor 488 (1:200, A-11055, Thermo Fisher Scientific), donkey anti-rat Alexa Fluor 488 (1:200, A-21208, Thermo Fisher Scientific), donkey anti-goat Alexa Fluor 555 (1:200, A-21432, Thermo Fisher), donkey anti-mouse Alexa Fluor 555 (1:200, A-31570, Thermo Fisher Scientific), chicken anti-rat Alexa Fluor 647 (1:200, A-21472, Thermo Fisher Scientific), donkey anti-rabbit Alexa Fluor 647 (1:200, A-31573, Thermo Fisher Scientific)) and DAPI (1 µg/mL, Merck) in blocking buffer for 4 h at 4 °C, and washed three times for 10 min at RT. Slides were mounted in ProLong™ Gold Antifade Mountant (Thermo Fisher Scientific) and imaged after 24-48 h.

#### Immunolabeling of E14 cells

After fixation and blocking, E14 cells were incubated with the following primary antibodies in blocking buffer overnight at 4 °C: rat anti-SOX2 (1:200, 14-9811-82, Thermo Fisher Scientific) and mouse anti-OCT4 (1:200, MA1-104, Thermo Fisher Scientific). The next day, cells were washed three times in washing solution for 5 min at RT and incubated with secondary antibodies (donkey anti-rat Alexa Fluor 488 (1:200, A-21208, Thermo Fisher Scientific) and donkey anti-mouse Alexa Fluor 555 (1:200, A-31570, Thermo Fisher Scientific)) and DAPI (1 µg/mL, Merck) in blocking buffer for 3 h at 4 °C. Finally, the cells were washed three times in washing solution for 5 min at RT and imaged directly.

#### Single-molecule fluorescence in-situ hybridization (smFISH)

smFISH was performed as described previously^61^. Briefly, samples were fixed with PFA and stored in 70% ethanol, as described above. Custom designed smFISH probes for *Dab2*, *Fst*, *Hhex* and *Spink1* (BioCat, Supplementary Table 5), labeled with Quasar 570, CAL Fluor Red 610, or Quasar 670, were incubated with the samples overnight at 30 °C in hybridization buffer (100 mg/mL dextran sulfate, 25% formamide, 2X SSC, 1 mg/mL E.coli tRNA, 1 mM vanadyl ribonucleoside complex, 0.25 mg/mL BSA; Thermo Fisher Scientific). Samples were washed twice for 30 min at 30 °C with wash buffer (25% formamide, 2X SSC). The wash buffer was supplemented with DAPI (1 μg/mL) in the second wash step. All solutions were prepared with RNAse-free water. Finally, the samples were mounted in ProlongGold (Life Technologies) and imaged when hardened (sections) or immediately (ibidi dishes). All components are from Sigma-Aldrich unless indicated.

#### Imaging

Fixed and stained samples were imaged on a Nikon Ti-Eclipse epifluorescence microscope equipped with an Andor iXON Ultra 888 EMCCD camera and dedicated, custom-made fluorescence filter sets (Nikon). Primarily, a 10× / 0.3 Plan Fluor DLL objective, a 20× / 0.5 Plan Fluor DLL objective, or a 40× / 1.3 Super Fluor oil-immersion objective (Nikon) were used. To image complete sections of cerebral organoids, multiple adjacent fields of view were acquired and combined using the tiling feature of the NIS Elements software (Nikon). Z-stacks were collected of whole-mount gastruloids and XEGs with distances of 10 μm between planes. For smFISH measurements, z-stacks were collected with a distance of 0.2 μm between planes in four fluorescence channels (DAPI, Quasar 570, CAL Fluor Red 610, Quasar 670) using a 100× /1.45 Plan Apo Lambda oil (Nikon) objective. Time lapses to observe the formation of epithelial structures were performed 24 h and 48 h after cell seeding, on XEGs grown from the mCherry-GPI ES cell line. XEGs were transferred to a glass-bottom μ-Slide imaging chamber (ibidi) and imaged every 30 min for 24 h with a Nikon Eclipse Ti C2+ confocal laser microscope (Nikon, Amsterdam, The Netherlands), equipped with lasers at wavelengths 408, 488 and 561, an automated stage and perfect focus system at 37°C and 5% CO_2_. Images were acquired with a Nikon 20x Dry Plan Apo VC NA 0.75 objective. To track SOX1 expression in gastruloids and XEGs during the 24 h growth after the GSK3 inhibitor pulse, 72 h gastruloids and XEGs grown from the *Sox1^GFPiresPac^* ES cell line were transferred to a glass-bottom μ-Slide imaging chamber (ibidi) and imaged every 40 min for 24 h, while temperature and CO_2_ levels were maintained at 37 °C and 5%, respectively, by a stage top incubator (INUG2-TIZW-SET, Tokai Hit) mounted on the Nikon Ti-Eclipse epifluorescence microscope.

#### Single-cell RNA-seq library preparation and sequencing

For each replicate, 96 pooled gastruloids and 96 pooled XEGs were collected from a round-bottomed low-adherence 96-well plate in 15 mL Falcon tubes and pelleted by gentle centrifugation (500 r.p.m. for 2 min). No final aggregate was excluded from the collection. After washing with cold PBS, samples were resuspended in N2B27. Cells were then dissociated by 5 min incubation in TrypLE (Gibco) and gentle trituration with a pipet, centrifuged and resuspended in 1 mL of cold N2B27. Cells were counted to determine cell number and viability. For the first replicate, ES-mCherry-GPI were spiked in at a frequency of 5%. For the second replicate, E14 cells were collected from culture dishes and incubated for 30 min at 4 °C with CITE-seq cell hashing^62^ antibody Ab_CD15 (1:200) (Biolegend). XEN*-eGFP* were collected from culture plates and incubated for 30 min at 4 °C with CITE-seq cell hashing antibody Ab_CD140 (1:200) (Biolegend). In the gastruloid sample, labeled E14 cells were spiked in at a frequency of 5%, whereas in the XEG sample labeled E14 and XEN-*eGFP* were spiked in, both at a frequency of 5%. High viability of the cells in all samples was confirmed before 10X library preparation. Single-cell RNA-seq libraries were prepared using the Chromium Single Cell 3’ Reagent Kit, Version 3 Chemistry (10x Genomics) according to the manufacturer’s protocol. CITE-seq libraries were prepared according to the CITE-seq protocol from New York Genome Center version 2019-02-13. Libraries were sequenced paired end on an Illumina Novaseq6000 at 150 base pairs.

## Computational methods

### Analysis of single-cell RNA-sequencing data

#### Single-cell RNA-seq data pruning and normalization

Cells with a low number of transcripts were excluded from further analysis based on the histograms in Supplementary Fig. 3a (count < 1300 for replicate 1 of the XEG experiment and count < 2300 for the other datasets). Genes expressed in less than 2 cells (across merged replicates) were excluded from further analysis. The final XEG dataset contains 14286 genes and 4591 or 6857 cells for replicate 1 or 2, respectively. The gastruloid dataset contains 14384 genes and 4233 or 8363 cells per replicate. The two datasets were normalized using the scran R-package (V 1.10.2^63^). Gene variabilities were calculated (improvedCV2, scran) for each replicate separately, after excluding ribosomal genes [Ribosomal Protein Gene Database, http://ribosome.med.miyazaki-u.ac.jp/], exogenously expressed genes and genes expressing the antibodies used for CITE-seq. The 10% most highly variable genes (HVG) were selected based on variability p-values.

#### Dimensionality reduction

For each of the two datasets, the two replicates were batch corrected with the fast mutual nearest neighbors (MNN) method implemented in the scran R-package^64^, using the union of the 10% HVG of the two replicates and log-transformed normalized counts with d = 120 (number of principal components) and k = 50 (number of nearest neighbours). For dimensionality reduction, a uniform manifold approximation and projection (UMAP) was calculated on the batch corrected data using the R-package UMAP (V 0.2.3.1) with n = 50, min_dist = 0.7 and using the cosine distance measure.

#### Identification of spike-in cells

Cells with any expression of mCherry were annotated as ES (mCherry+). The remaining spike-in cells, E14 (CD15+) and XEN spike-in (CD140+) (see Single-cell RNA-seq library preparation and sequencing), could not be determined by the expression level of the antibody alone. We therefore chose to assign spike-ins based on clusters. For each of the two datasets, a shared nearest neighbor graph was constructed from the batch corrected data (see Dimensionality reduction) with scran using k = 20 and d = 30. Louvain clustering was performed on the constructed graphs with the R-package igraph (V1.2.4.1), which resulted in 8 clusters for XEGs and 7 clusters for gastruloids (see Supplementary Fig. 3c). We identified 3 out of the 8 clusters in XEGs based on literature markers and spike-in gene expression. One cluster out of these three was mainly comprised of mESCs, due to high Ab_CD15 expression and mCherry positive cells. Cells that had an expression of Ab_CD15 > 50 and were part of this cluster were considered spiked-in E14 and annotated as E14 (CD15+). The other two clusters were both eGFP positive, where one of them had a higher Ab_CD140 expression and was thus annotated as XEN spike-in (Ab_CD140+). The second cluster was annotated as XEN derived (Ab_CD140-). Similarly, for gastruloids, one of the 7 clusters was comprised of mainly mESCs based on literature markers and spike-in gene expression. Cells that had an expression of Ab_CD15 > 100 and were part of this cluster were considered spiked-in E14 and annotated as E14 (CD15+).

#### Analysis of cell cycle and stress-related genes

For each of the two datasets, cell cycle analysis was performed with the scran package using the cyclone function^65^ on the normalized counts. Cells in G2M phase were distributed evenly across all clusters and thus the clustering was not biased by cell cycle. No other separate cluster that consisted entirely of cell cycle related cells appeared.

For the analysis of stress-related genes, a list of known stress genes^66^ was used to calculate the average standardized expression per cell based on normalized counts. Stress-related genes were mainly found within the spike-in cells and there was no other separate cluster that consisted entirely of highly stressed cells.

#### Mapping to *in vivo* datasets

Our datasets were mapped to three different *in vivo* datasets.

##### Pijuan-Sala et al. dataset

The Pijuan-Sala et al. dataset^36^, which was downloaded from https://content.cruk.cam.ac.uk/jmlab/atlas_data.tar.gz, consists of 9 timepoints from E6.5 to E8.5. The data was normalized by size factors provided by the authors. Cells with no cell type assignment were excluded from further analysis. The 10% HVG were calculated (improvedCV2, scran package) on the remaining cells excluding sex genes, similar to Pijuan-Sala et al.’s method. Cells in the “mixed_gastrulation” cluster were also excluded. MNN mapping was applied to log-transformed normalized counts of the 10% HVG. First, *in vivo* timepoints were mapped to each other in decreasing order. Then, each of our four datasets was mapped separately to the combined Pijuan-Sala et al. dataset (MNN method with d = 120, k = 50). K-nearest-neighbor (knn) assignment was performed in the batch corrected principal component space. For each cell in our datasets, the 50 nearest neighbors in the *in vivo* dataset, based on Euclidean distances, were calculated. Each cell was assigned the most abundant cell type within the knn, if certain distance and confidence score conditions were met. This confidence score was calculated for each cell as the number of the most abundant cell type divided by the total number of neighbors (k=50). A cell was annotated as “Not assigned” if either, the average distance to its nearest neighbor exceeded a certain threshold (determined by the long tail of the histogram of average distances for each of our datasets separately) or the assignment had a confidence score less than 0.5. Additionally, we placed cells in “Not assigned” if they were assigned to clusters with less than 10 cells, or to the cluster “Blood progenitors 2” (because this cluster did not show distinct expression of known literature markers). This resulted in 22 assigned clusters for XEGs and 15 assigned clusters for gastruloids. For each cell in our dataset we calculated the average and the standard deviation of the developmental age of the knn.

##### Nowotschin et al. dataset

The Nowotschin et al. dataset^41^, which was downloaded from https://endoderm-explorer.com/, consists of 6 timepoints from E3.5 to E8.75. The data was normalized (scran) and the 10% HVG were calculated (improvedCV2, scran package). First, MNN was applied to the Nowotschin et al. dataset in increasing order of the timepoints (using log-transformed normalized counts of the 10% HVG, d = 150, k =50). Then, XEN cells from our XEG dataset (XEN spike-ins (CD140+) and XEN derived (CD140-)) were mapped to the MNN-corrected Nowotschin et al. dataset. Knn assignment was performed as described above and resulted in 7 assigned clusters.

##### Delile et al. dataset

The Delile et al. dataset^37^, which was downloaded from https://github.com/juliendelile/MouseSpinalCordAtlas, consists of 5 timepoints from E9.5 to E13.5. Cells that had a cell type assignment of “Null” or “Outlier” were excluded from further analysis. The data was normalized (scran) and the 10% HVG were calculated. First, MNN was applied to the Delile et al. dataset in order of increasing timepoints (log-transformed normalized counts of the 10% HVG, d = 120, k =50). Then, we mapped “Spinal cord” cells to the MNN corrected Delile et al. dataset separately for each of our replicates. Knn assignment was performed as described above and resulted in 3 clusters for XEGs and 3 clusters for gastruloids.

#### Differential expression analysis

For the differential expression test between “spike-in XENs” and “XENs in XEGs” a Welch t-test (implemented in findMarkers, scran R package) was conducted on the normalized log-transformed counts. The test was performed on XEGs from replicate 2. “spike-in XENs” were chosen as the 100 cells with highest Ab_CD140 expression and “XENs in XEGs” were the 100 cells with lowest Ab_CD140 expression within the XEN identified cells.

For the differential expression test between XEGs and gastruloids, a negative binomial regression was performed (R package edgeR V 3.24.3^67^). Based on the knn assignment to the Pijuan-Sala et al. dataset, all cells annotated as “Spinal cord” were extracted from our four datasets (in XEGs 859 cells in replicate 1 and 166 cells in replicate 2, in gastruloids 2071 cells in replicate 1 and 1882 cells in replicate 2). Raw counts were used for the regression with these four subsets as dummy variables and a variable corresponding to the total number of counts per cell. P-values were obtained for the contrast between XEGs and gastruloids using the average regression coefficients among variables of both replicates.

Similarly, for the differential expression test of the “Spinal cord” in XEGs, a negative binomial regression was used. Cells were excluded from the test if either their cell type occurred in less than 10 cells per replicate, or if the cells were annotated as “Not assigned”, leaving a total of 13 cell types (7742 cells) to be considered. For each cell type and each replicate a dummy variable was created and a variable corresponding to the total number of counts per cell. Then, p-values were obtained for the contrast between the average regression coefficients of the two replicates of the “Spinal cord” cluster and the average regression coefficients of all other variables considered in the test.

For all differential expression tests p-values were adjusted for multiple hypothesis testing with the Benjamini-Hochberg method.

#### Scatter plot of dorsoventral markers

For the scatter plot of canonical markers, we used the average log-transformed normalized expression of 10 canonical dorsal markers (*Pax7, Msx3, Pax3, Gsh2, Msx2, Olig3, Math1, Zic1, Msx1, Zic2*) and the average log-transformed normalized expression of 10 canonical ventral markers (*Olig2, Nkx6.1, Nkx2.2, Foxa2, Ncam1, Nrcam, Nkx6.2, Isl1, Fabp7, Hb9*). The solid line is a running average, calculated by sorting the data on ventral marker expression and averaging dorsal marker expression in sliding bins of 50 data points. The scatterplot of Msx1 and Nrcam module expression was calculated similarly, using 10 genes with the highest Spearman correlation with Msx1 or Nrcam, respectively.

### Image analysis

Image stacks of whole-mount immunostained gastruloids and XEGs, and images of immunostained sections were pre-processed by background subtraction (rolling ball, radius: 50 pixels = 65 μm (10× objective), 32 μm (20× objective) or 16 μm (40× objective)) in the channels that showed autofluorescent background using ImageJ^68^. When background subtraction in images of sections did not result in proper removal of autofluorescent background signal, the Enhance Local Contrast (CLAHE) tool was used in ImageJ^68^. smFISH image stacks were pre-processed by applying a Laplacian of a Gaussian filter (σ = 1) over the smFISH channels using scikit-image (v0.16.1) ^69^. For all image stacks, a maximum projection was used to obtain a 2D representation. To show a single object per image, images were cropped around the object of interest.

## Acknowledgements

We are thankful to Alfonso Martinez Arias for insightful discussions and feedback on the manuscript. We acknowledge Anna-Katerina Hadjantonakis for helpful input at various stages of the project. We also thank Dr. Sylvia Le Dévédec and Hans de Bont of the Leiden University Cell Observatory for their support in this work. N. B.-C., M. M., P. v.d. B., M. F. and S.S. were supported by the Netherlands Organisation for Scientific Research (NWO/OCW, www.nwo.nl), as part of the Frontiers of Nanoscience (NanoFront) program. E.A. acknowledges support by a Stichting voor Fundamenteel Onderzoek der Materie (FOM, www.nwo.nl) projectruimte grant (16PR1040). M.H. acknowledges support by a Netherlands Organisation for Scientific Research (NWO/OCW, www.nwo.nl) VIDI grant (016.Vidi.189.007). This work was carried out on the Dutch national e-infrastructure with the support of SURF Cooperative. The funders had no role in study design, data collection and analysis, decision to publish, or preparation of the manuscript.

## Author contributions

N.B.-C., E.A. and M.H. cultured gastruloids and XEGs. N.B-C., E.A. and M.H. performed signaling experiments and immunostaining and analyzed the resulting images, N.B-C. prepared samples for single-cell RNA sequencing and interpreted the sequencing data, M.M. performed the computational analysis of the single-cell RNA sequencing data, P.v.d.B. contributed to the computational analysis of single-cell RNA sequencing data and carried out the smFISH measurements, M.F. supported all experiments and performed all cryosectioning, N.B.-C., M. M., E.A., P.v.d.B. and M.H. produced figures, N.B.-C., M. M., E.A., P.v.d.B., M.H. and M.F. contributed to the manuscript, T.I., S.T. and S.S. conceived the study and acquired funding. S.S. interpreted the data and wrote the manuscript. All authors discussed the results and commented on the manuscript at all stages.

## Data availability

The single-cell RNA sequencing datasets generated in this study are available in the Gene Expression Omnibus repository, <u>GSE141530</u>.

## Code availability

Custom R and python code used to analyze the data is available from the authors upon request.

## Correspondence and requests for materials

Should be addressed to S.S., S.T., T.I., or M.H.

---

**Supplementary Fig. 1.**
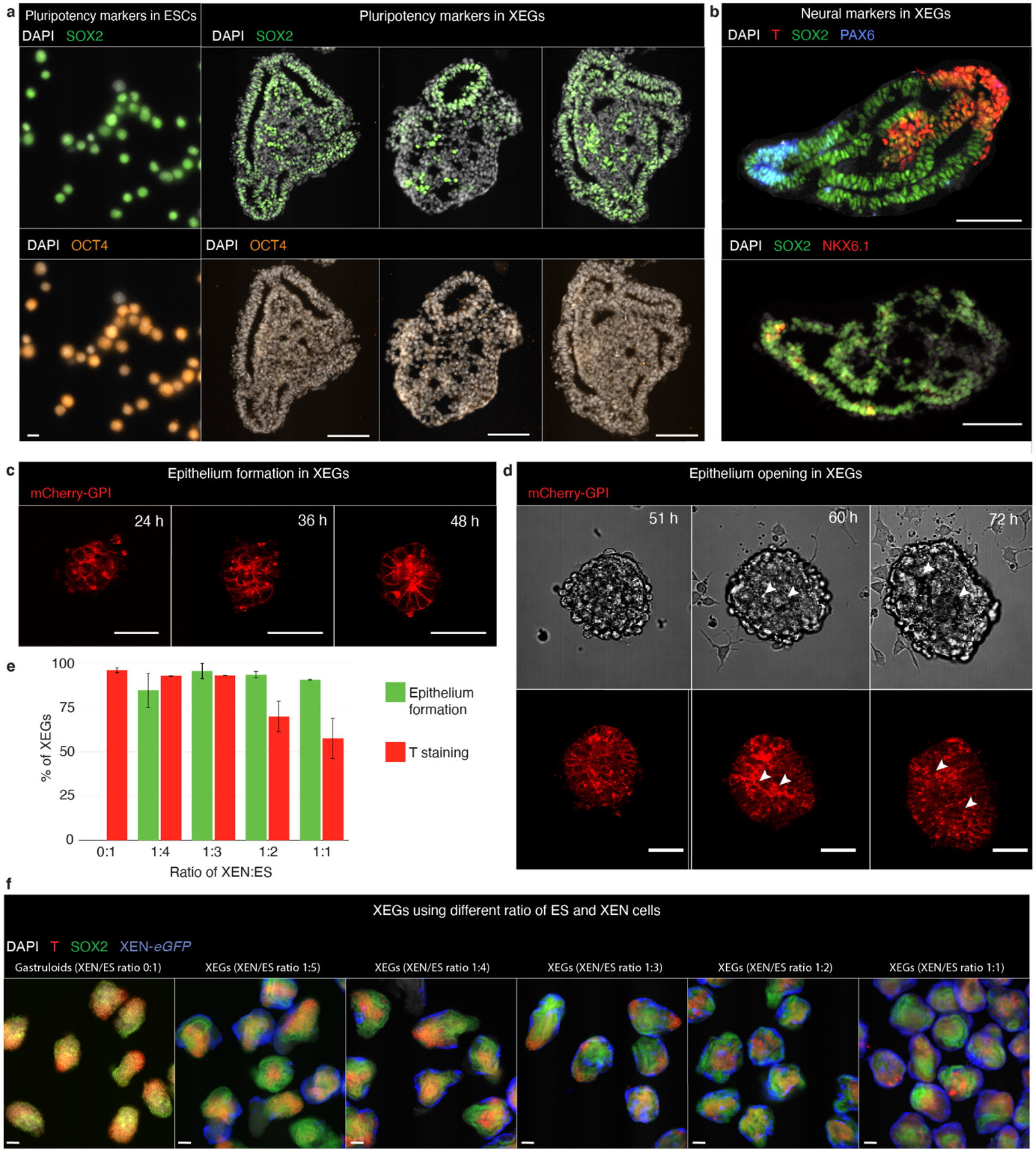
Optimization and characterization of XEGs. **a**, Expression of SOX2 and OCT4 (immunostaining) in sections of XEGs at 96 h (left, scale bars: 100 µm) and cultured ESCs (right, scale bars: 10 µm). **b**, Expression of T, SOX2, PAX6, MASH1 and NKX6.1 (immunostaining) in 96 h XEGs. Scale bars: 100 µm. **c-d,** Live-cell imaging of morphological changes in XEGs grown from mCherry-GPI expressing mESCs. mCherry-GPI is localized to the cell membrane. In all images, a single z-plane is shown. **c**, Rosette formation. The center of the rosette is indicated by a dashed line. Inset: tracing of cell outlines. **d**, Cavitation of rosettes. Dashed lines indicate the opening cavities. The top row shows the brightfield channel, the bottom row shows the mCherry channel. See also Supplementary Videos 1 and 2. Scale bars: 50 µm. **e**, Average fraction of aggregates showing epithelial structures and T staining at 96 h for different starting ratios of ESCs and XEN cells (n = 2 experiments, error bars show standard deviation). **f**, SOX2 (neural progenitors-like cells) and T (primitive streak-like cells) expression in gastruloids and XEGs with different starting ratios of ESCs and XEN cells (z-projection of whole mount immunostaining). Scale bars: 100 µm. **a, b, f,** Cell nuclei were stained with DAPI.

**Supplementary Fig. 2.**
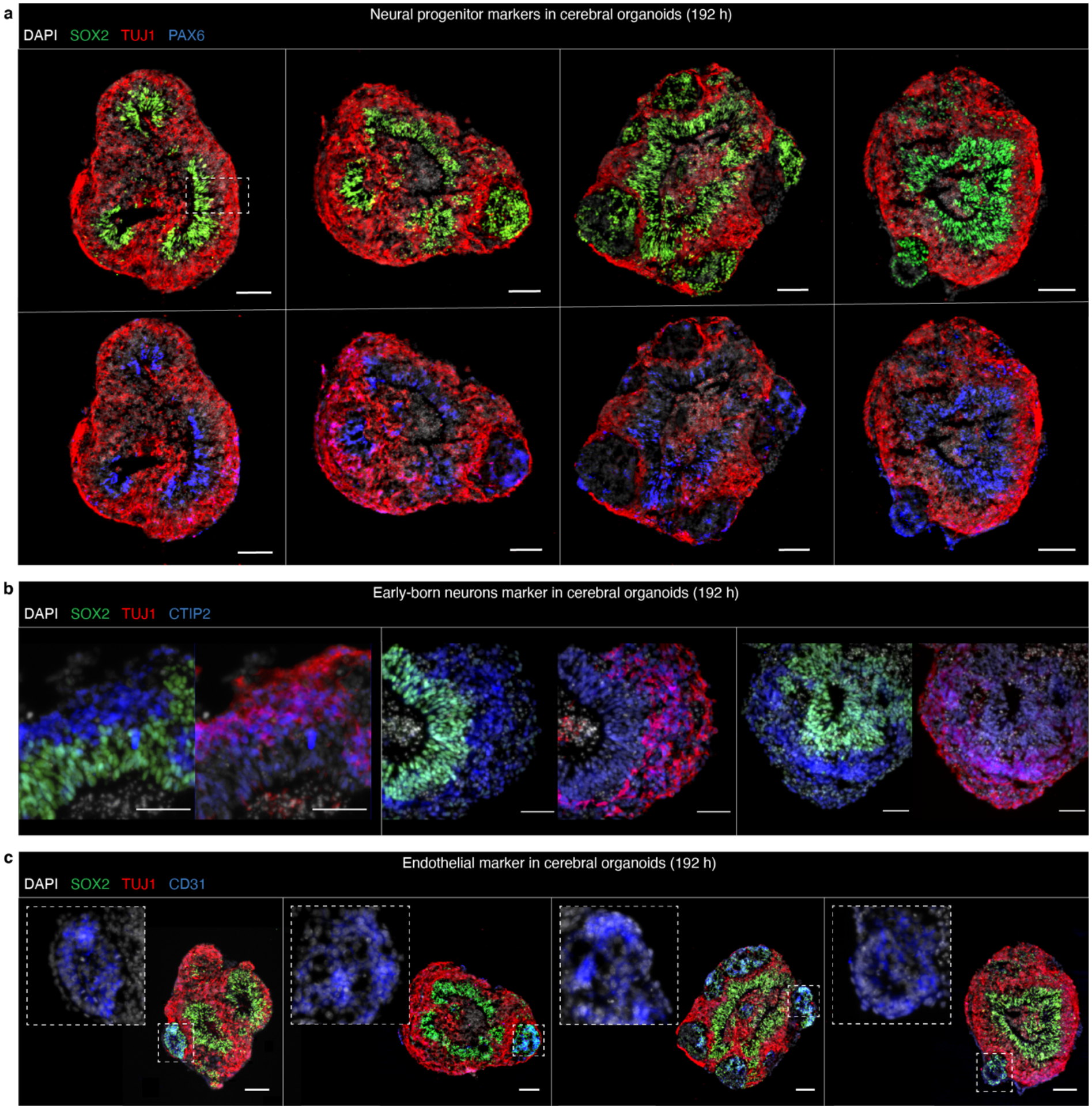
Developing brain markers are expressed in XEGs further differentiated with a cerebral organoid protocol. **a, b, c,** Immunostaining in sections of XEGs on day 8 after cell seeding. **a**, TUJ1, SOX2 (top) and PAX6 (bottom). The dashed box highlights an example of layered organization adjacent to a ventricle-like cavity. **b,** TUJ1, SOX2 and CTIP2, a marker for early-born neurons. **c,** TUJ1, SOX2 and CD31. Insets show clusters of cells positive for the endothelial marker CD31. Cell nuclei were stained with DAPI. Scale bars: 100 µm.

**Supplementary Fig. 3.**
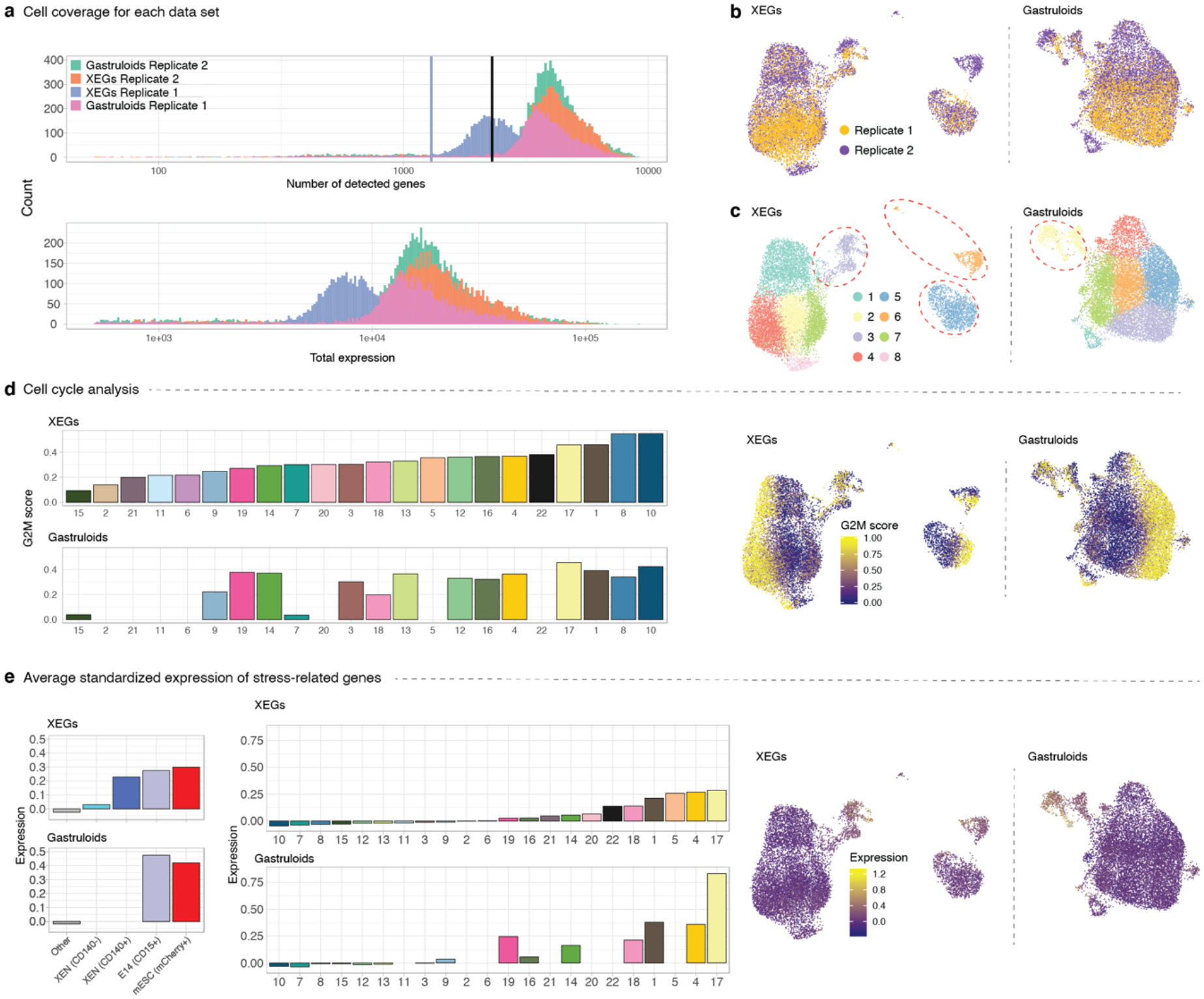
Quality control of single-cell RNA-seq data. **a**, Top, number of detected genes per cell in each replicate; the blue line indicates a quality control threshold for XEGs from replicate 1 and the black line for the remaining datasets. Bottom, total expression per cell for each dataset. **b**, UMAP of cells in XEGs and gastruloids, colored by replicate. **c**, UMAP of cells in XEGs and gastruloids, colored by Louvain clustering. The encircled clusters contain the spiked-in cells. **d**, Left, average G2M scores for each cell type. Right, UMAPs of cells in XEGs and gastruloids colored by G2M score. **e**, Left, average standardized expression of stress-related genes in spike-in cells. Middle, expression of stress-related genes by cell type. Right, UMAPs of cells in XEGs and gastruloids with expression of stress-related genes indicated by color. **b-e**, UMAPs contain both replicates, batch corrected.

**Supplementary Fig. 4.**
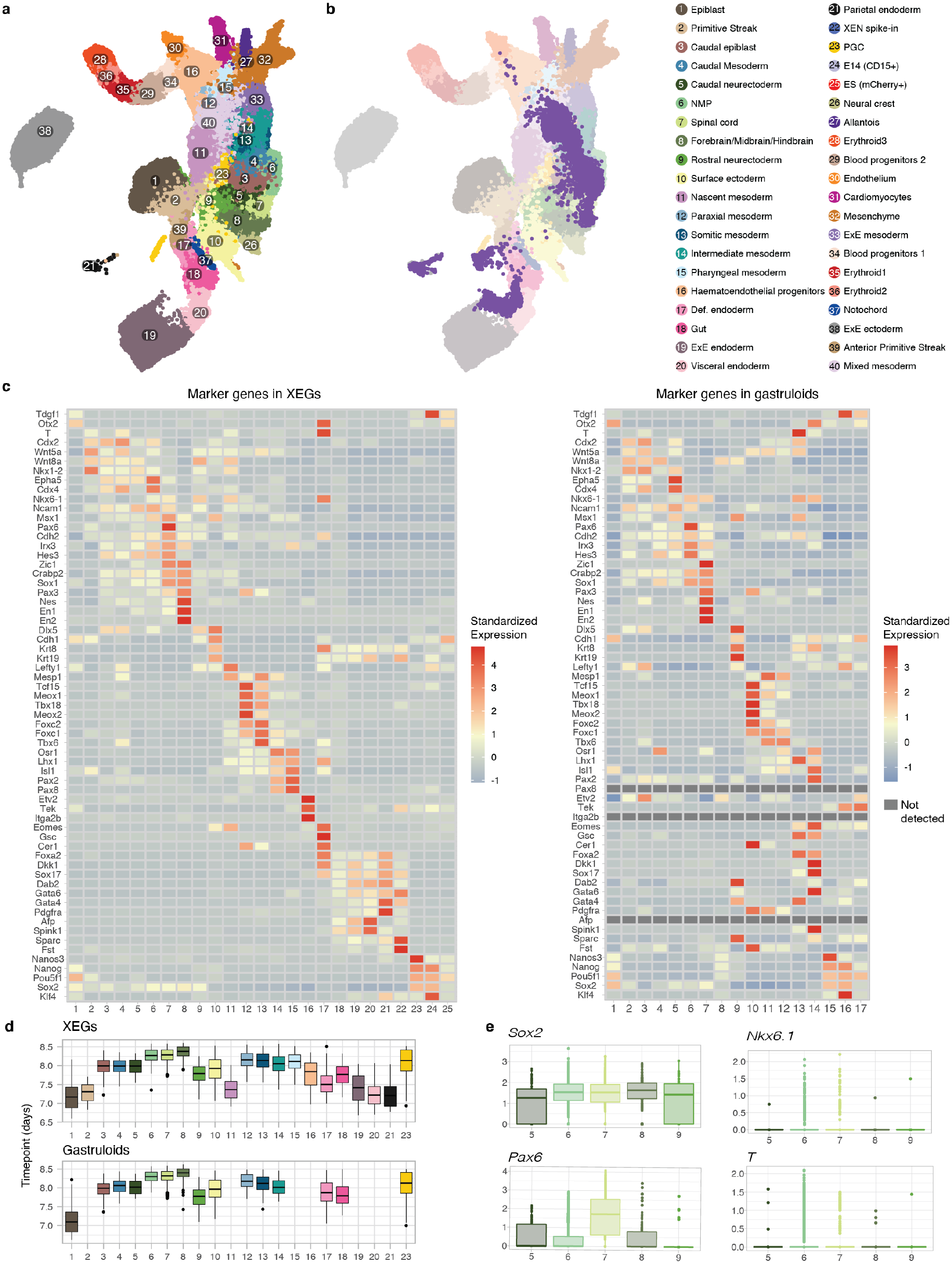
Single-cell RNA-seq resolves cell type diversity in XEGs and gastruloids. **a**, UMAP of the Pijuan-Sala et al.^36^ dataset with cell types indicated by color. **b**, MNN mapping of XEG cells from replicate 2 (bright colors) to the Pijuan-Sala et al. dataset (dim colors), as an example for the mapping procedure. **c**, Heat map of standardized expression of genes associated with mouse embryonic development in XEGs and gastruloids. References describing the *in vivo* expression of the genes are given in Supplementary Table 1. **d**, Developmental age of cell types based on mapping to *in vivo* data. **e**, Expression of *Sox2*, *Pax6* and *Nkx6.1* and *T* in XEGs, as measured by single-cell RNA-seq.

**Supplementary Fig. 5.**
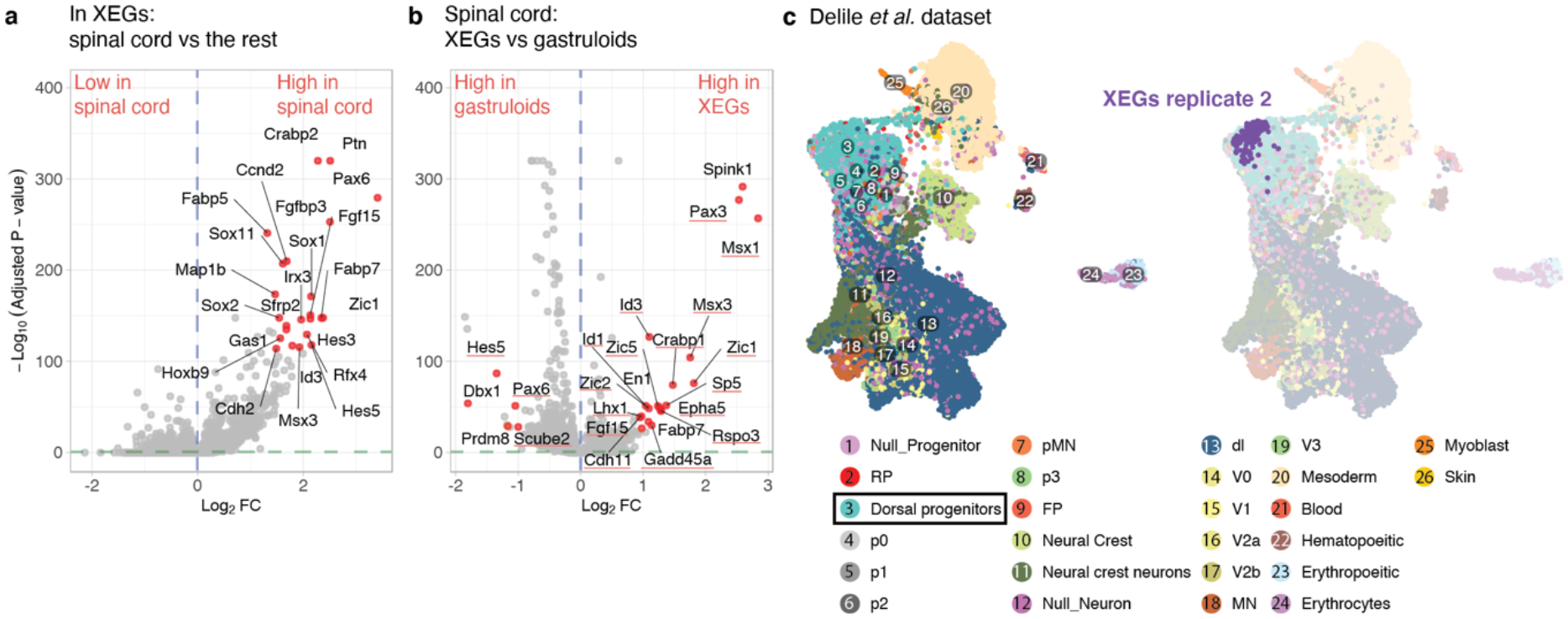
Neuroepithelial cells are biased towards a dorsal expression profile. **a**, Gene expression differences between cells classified as “spinal cord” and all other cells in XEGs (fold-change vs p-value). Named genes are expressed in the neural tube according to previous studies (Supplementary Table 2). **b**, Gene expression differences between cells classified as “spinal cord” in gastruloids and XEGs (fold-change vs p-value). Underlined genes are expressed in the dorsal part of the neural tube according to previous studies (Supplementary Table 4). **c**, Left, UMAP of the cells in the Delile et al. dataset^37^, colored by cell type. Right, MNN mapping of cells classified as “spinal cord” in replicate 2 XEGs (bright colors) to the Delile et al. dataset (dim colors), as an example of the mapping procedure.

**Supplementary Fig. 6.**
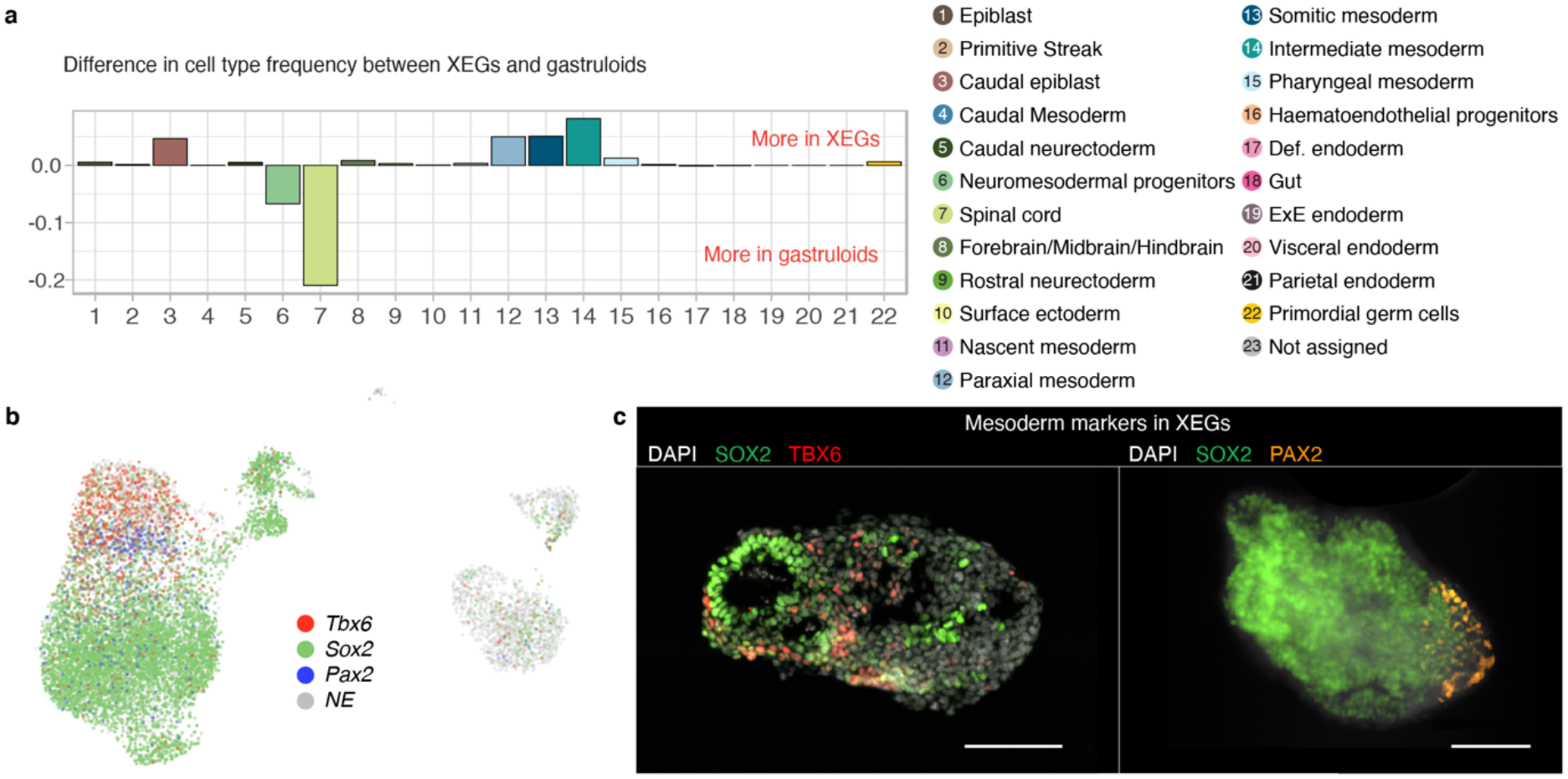
XEGs and gastruloids differ in cell type distribution. **a,** Differences between relative frequencies of cell types in XEGs and gastruloids. **b,** UMAP of cells in XEGs (both replicates, batch corrected) with log expression of *Tbx6*, *Sox2* and *Pax2* indicated by color. **c,** Expression of mesoderm markers. Left, TBX6 expression in a 96 h XEG (immunostaining of a section). Right, PAX2 expression in a 96 h XEG (wholemount immunostaining). Scale bars: 100 µm. Nuclei were stained with DAPI.

**Supplementary Fig. 7.**
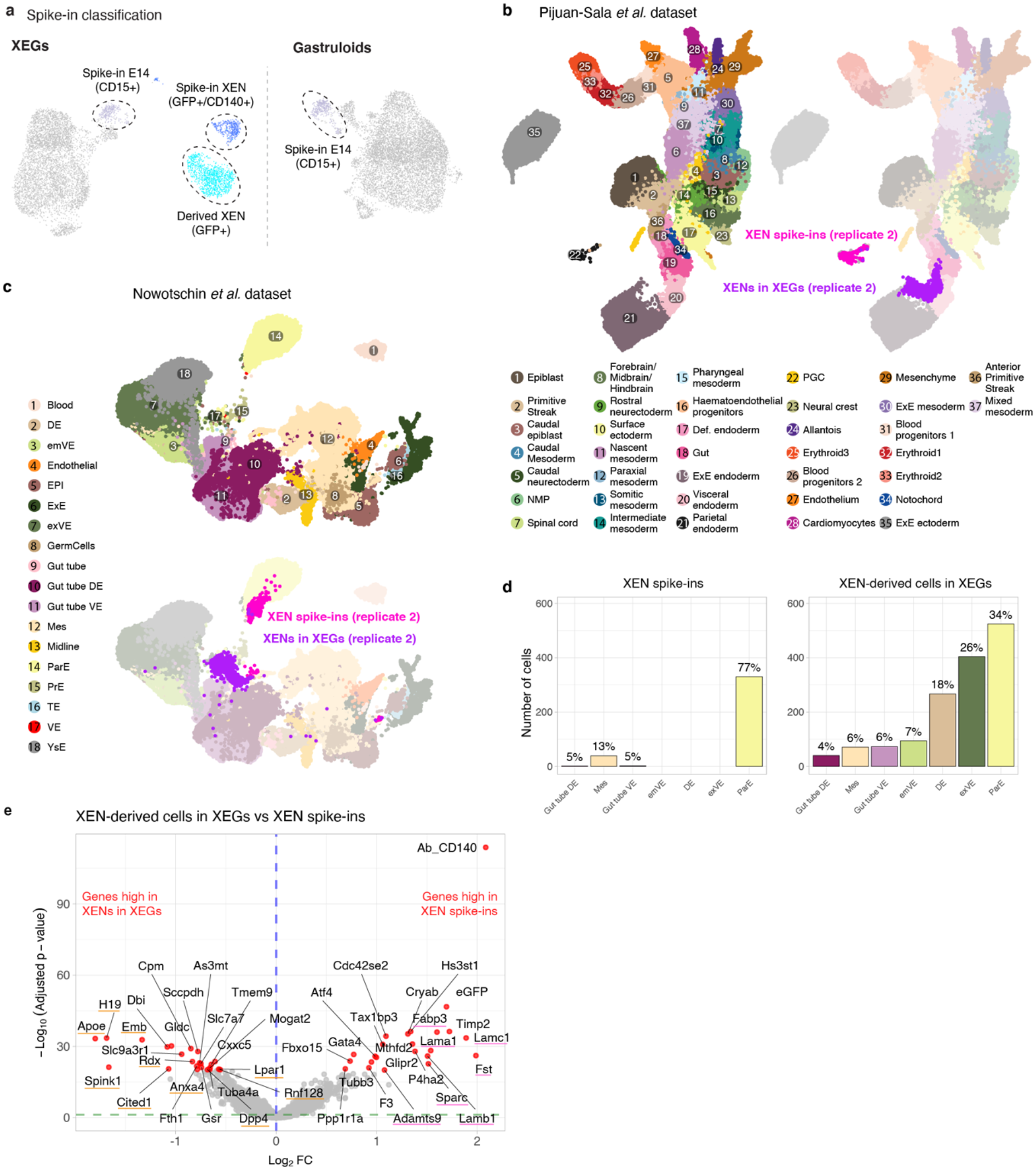
Most XEN-derived cells are visceral endoderm-like in XEGs. **a**, UMAP of cells in XEGs and gastruloids with spike-in cells and XEN derived cells highlighted by color and circled (replicate 2). **b**, **c**, UMAPs of the Pijuan-Sala^36^ or Nowotschin^41^ dataset, respectively. XEN spike-ins and XEN-derived cells from XEGs replicate 2 (bright colors) are mapped to the *in vivo* datasets (dim colors). **d**, Cell type frequencies of XEN spike-ins and XEN derived cells in XEGs, resulting from knn assignments based on the mapping in **(c)**. **e**, Gene expression differences between XEN spike-ins and XEN-derived cells in XEGs (fold-change vs p-value). Orange and pink lines indicate genes with PE-like and VE-like identity, respectively (see Supplementary Table 3). DE: definitive endoderm, emVE: embryonic visceral endoderm, EPI: epiblast, ExE: extraembryonic ectoderm, exVE: extraembryonic visceral endoderm, Mes: mesoderm, ParE: parietal endoderm, PrE: primitive endoderm, TE: trophectoderm, VE: visceral endoderm, YsE: yolk sac endoderm.

**Supplementary Fig. 8.**
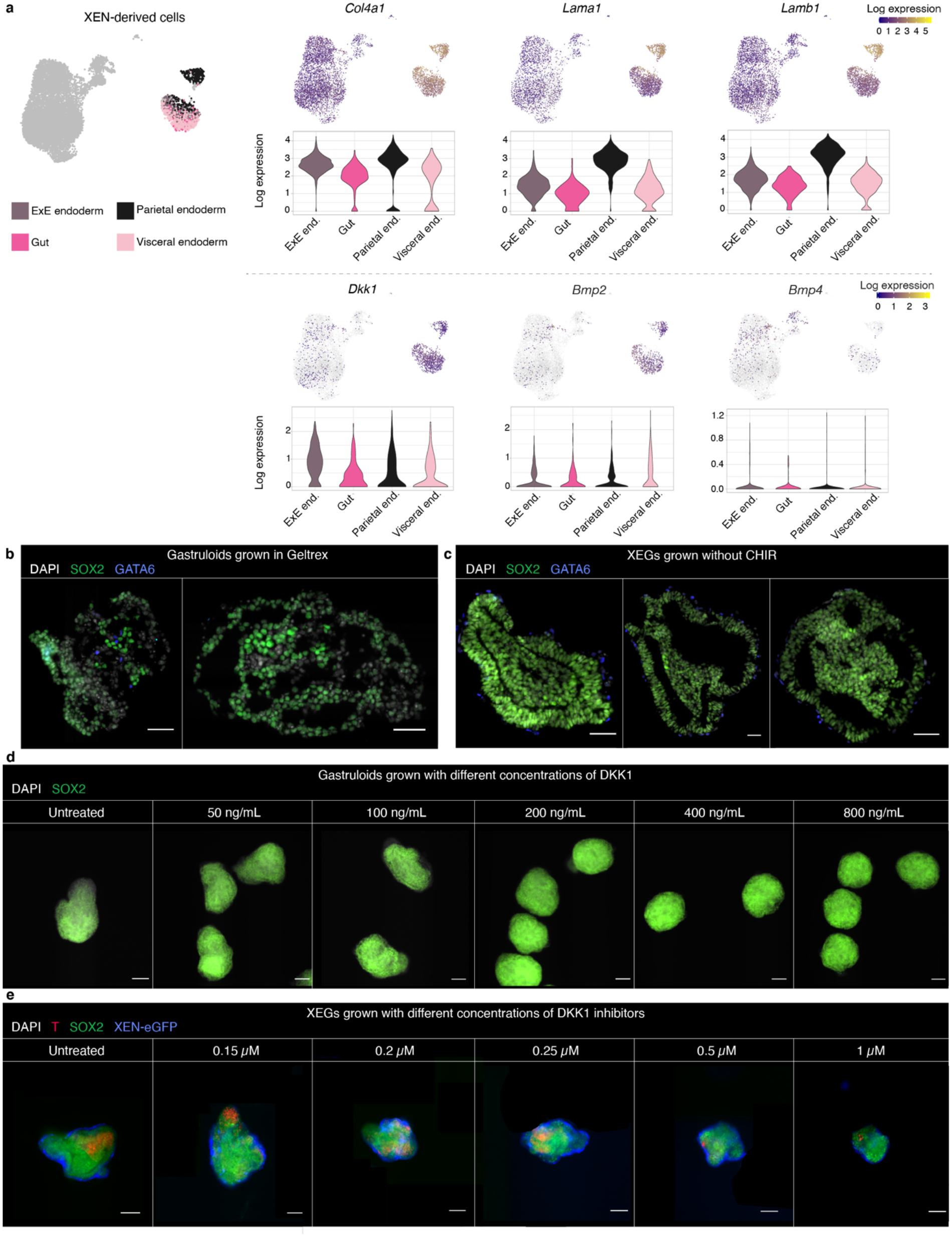
Basement membrane components and the WNT inhibitor DKK1, all expressed exclusively by XEN cells, play a role in epithelia formation. **a**, Expression of genes in XEN-derived cells. Left, UMAP of cells in XEGs with XEN-derived cells colored by cell type (gut, parietal endoderm (parietal end.), embryonic VE (visceral end.) or extraembryonic VE (ExE end.)). Top-right, expression of basement membrane components collagen IV *(Col4a1)*, laminin alpha 1 (*Lama1*) and laminin beta 1 (*Lamb1*). Bottom-right, expression of signaling factors DKK1 (*Dkk1*), BMP2 (*Bmp2*) and BMP4 (*Bmp4*). UMAPs indicate log expression by color and contain both replicates, batch corrected. A violin plot of log expression in XEN-derived cell types is shown below the UMAP for each gene. **b**, Expression of SOX2 and GATA6 in gastruloids grown in Geltrex at 96 h (immunostaining of crysections). Scale bars: 50 µm. **c**, Expression of SOX2 in XEGs grown without CHIR (immunostaining of cryosections). No specific T staining could be detected (data not shown). XEN cells were localized by expression of GATA6. Scale bars: 50 µm. **d**, Expression of SOX2 in 96 h gastruloids treated with various concentrations of DKK1 between 24 h and 96 h (wholemount immunostaining). Scale bars: 100 µm. **e**, Expression of SOX2 and T in 96 h XEGs treated with various concentrations of DKK1 inhibitor WAY-262611 between 24 h and 96 h (wholemount immunostaining). XEN cells were localized by endogenous expression of H2B-GFP. Scale bars: 100 µm. **b**-**e**, Cell nuclei were stained with DAPI.

**Supplementary Table 1.**
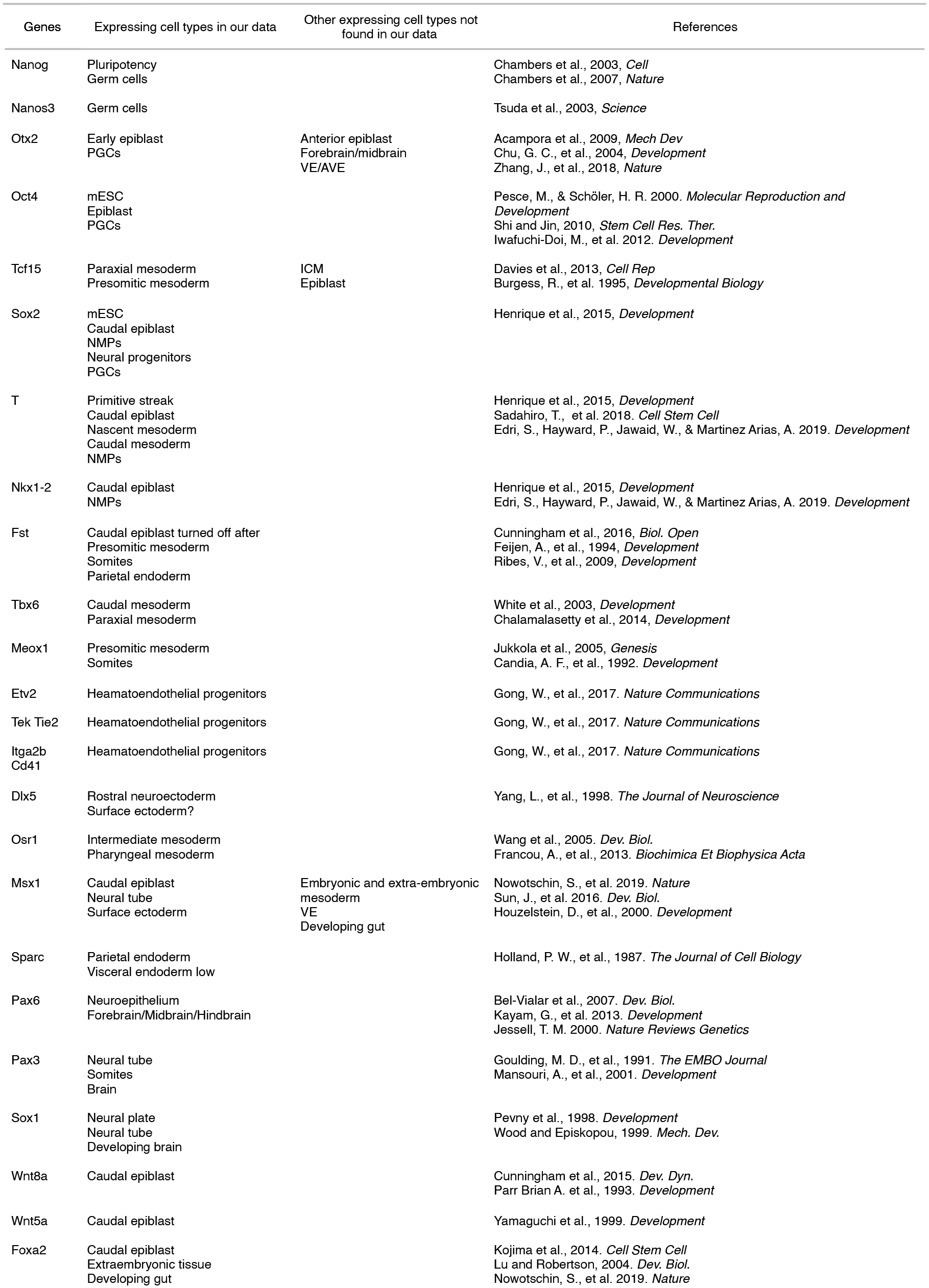
Marker genes used for cell type annotation in the single-cell RNA-seq data.

**Supplementary Table 2.** Genes differentially expressed between spinal cord-like cells and the other cells types in XEGs.

| Gene | Log FC | p.adjust | Reference |
| --- | --- | --- | --- |
| Ccnd2 | 1.684190185 | 1.37E-210 | Lacomme, M., et al., (2012). *Molecular and Cellular Biology* |
| Crabp2 | 2.27328271 | 0 | Ruberte, E., et al., (1992). *Development* |
| Id3 | 1.789970808 | 5.65E-118 | Wine-Lee, L., et al., (2004). *Development* |
| Fabp7 (BLBP) | 2.339513279 | 2.65E-148 | Feng, L., Hatten, M. E., & Heintz, N. (1994). *Neuron* |
| Rfx4 | 2.066843727 | 3.40E-130 | Ashique, A. M., et al. (2009). *Science Signaling* |
| Hoxb9 | 1.561341789 | 4.07E-126 | Graham, A., Maden, M., & Krumlauf, R. (1991). *Development* |
| Cdh2 | 1.486458795 | 1.15E-114 | Radice, G. L., et al., (1997). *Developmental Biology* |
| Msx3 | 1.925035016 | 3.06E-116 | Misra, K., Luo, H., Li, S., Matise, M., & Xiang, M. (2014). *Development* |
| Pax6 | 3.403368474 | 4.04E-280 | Jeong, J., & McMahon, A. P. (2005). *Development* |
| Fabp5 | 1.316127675 | 1.97E-241 | Shimamoto, C., et al. (2014). *Human Molecular Genetics* |
| Sfrp2 | 1.677002264 | 9.74E-140 | Misra, K., & Matise, M. P. (2010). *Developmental Biology* |
| Hes3 | 2.132129411 | 2.12E-147 | Lobe, C. G. (1997). *Mechanisms of Development* |
| Ptn | 2.50622717 | 0 | Magdaleno, S., et al. (2006). *PLoS Biology* |
| Fgf15 | 2.131102328 | 2.17E-151 | McWhirter, J. R., et al., (1997). *Development* |
| Irx3 | 1.957663655 | 1.71E-146 | Lebel, M., et al. (2003). *Molecular and Cellular Biology* |
| Zic1 | 2.371434543 | 1.56E-148 | Nagai, T., et al., (1997). *Developmental Biology* |
| Fgfbp3 | 2.502793725 | 1.10E-253 | Yu, K., McGlynn, S., & Matise, M. P. (2013). *Development* |
| Hes5 | 2.152139781 | 9.35E-119 | Hatakeyama, J., et al., (2004). *Development* |
| Map1b | 1.46443751 | 3.03E-174 | Fawcett, J. W., et al., (1994). *Neuroscience* |
| Gas1 | 1.676359071 | 1.03E-135 | Allen, B. L., Tenzen, T., & McMahon, A. P. (2007). *Genes & Development* |
| Sox11 | 1.610646468 | 6.85E-208 | Tanaka, S., et al., (2004). *Molecular and Cellular Biology* |
| Sox2 | 1.54213575 | 1.78E-148 | Hoffmann, S. A., et al., (2014). *Development* |
| Sox1 | 2.143957789 | 5.90E-172 | Hoffmann, S. A., et al., (2014). *Development* |

**Supplementary Table 3.** Genes differentially expressed between XEN-derived cells in XEGs and cultured XEN cells.

| Gene | Log FC | Expression summary | Reference |
| --- | --- | --- | --- |
| Apoe | -1.80 | VE > PE | Basheeruddin, K., et al., (1987) |
| H19 | -1.69 | VE > PE | Long, L., & Spear, B. T. (2004). |
| Spink1 | -1.66 | VE | Goh, H. N., et al. (2014). |
| Emb | -1.33 | pan-endoderm | Brown, K., et al., (2010). |
| Cited1 (Mgs1) | -1.07 | exVE | Dunwoodie, S. L., et al., (1998). |
| Rdx (Radixin) | -0.83 | VE | Igarashi, H., et al. (2018). McClatchey, A. I., et al.,(1997). |
| Anxa4 | -0.78 | pan-endoderm | Brown, K., et al., (2010). |
| Dpp4 | -0.68 | VE | Sherwood, R. I., et al. (2007). |
| Laprl (Lpa1) | -0.57 | VE | Koike, S., et al., (2009). |
| Rnf128 (Greul) | -0.56 | VE | Borchers, A. G. M., et al., (2002). |
| Gata4 | 0.77 | pan-endoderm | Morrissey, E. E., et al.,(1998). |
| P4ha2 | 0.79 | PrE | Ohnishi, Y., et al. (2014). |
| Adamts9 | 1.08 | PE | Jungers, K. A., et al., (2005). |
| Lamb1 | 1.51 | PE | Miner, J. H., et al., (2004). |
| Sparc | 1.51 | PE | Mason, I. J., et al., (1986). |
| Lama1 | 1.54 | PE | Hogan, B. L., et al., (1980). |
| Fabp3 | 1.60 | exVE - PE | Cheng, S., et al. (2019). Futaki, S., et al. (2003). |
| Lamc1 | 1.89 | PE | Smyth, N., et al., (1999). |
| Fst | 1.99 | PE | Feijen, A., et al. (1994). |

**Supplementary Table 4.** Genes differentially expressed between spinal cord-like cells in XEGs and gastruloids.

| Gene | Log FC | Expression summary | BMP signalling response | Reference |
| --- | --- | --- | --- | --- |
| Msx1 | 2.84 | Dorsal neural tube<br>+ small ventral domain | BMP (Lee, K. J., & Jessell, T. M. (1999)) | Misra, K., et al., (2014).<br>Duval, N., et al. (2014). |
| Spink1 | 2.59 |  |  |  |
| Pax3 | 2.53 | Dorsal neural tube | BMP (Lee, K. J., & Jessell, T. M. (1999)) | Kriks. S., et al., (2005). |
| Zic1 | 1.81 | Dorsal neural tube | BMP (Lee, K. J., & Jessell, T. M. (1999)) | Nagai, T., et al., (1997).<br>Aruga, J., et al., (2002). |
| Msx3 | 1.75 | Dorsal neural tube | BMP4 (Shimeld, S. et al., (1996)) | Misra, K., et al., (2014).<br>Duval, N., et al. (2014). |
| Crabp1 | 1.47 | Forebrain, Neural crest + dorsal neural tube | - | Ruberte, E., et al., (1991). |
| Sp5 | 1.37 | Midbrain and dorsal neural tube | - | Andoniadou, C. L., et al. (2011).<br>Dunty, W. C., et al., (2014). |
| Epha5 | 1.29 | Dorsal and ventral | BMP2 (Yamada, T., et al., (2016)) | Abdul-Aziz, N. M., et al., (2009).<br>Ono, K., et al. (2014). |
| Rspo3 | 1.28 | Forebrain, dorsal neural tube | - | Nam, J.-S., et al., (2007). |
| En1 | 1.27 | Hinbrain/Midbrain<br>Ventral neural tube | Not affected by BMP (Timmer, J. R., et al., (2002)) | Ericson, J., et al. (1997).<br>Sgaier, S. K., et al., (2007). |
| Zic5 | 1.23 | Dorsal neural tube and head fold | BMP (Nakata, K., et a., (2000)) | Inoue, T., et al., (2004). |
| Fabp7 (BLBP) | 1.14 | Ventral neural tube<br>Midbrain | - | Delille, J., et al., (2019).<br>Feng, L., et al., (1994). |
| Id3 | 1.10 | The roof plate and dorsal neural ectoderm<br>+ small ventral domain | BMP (Wine-Lee, L., et al., (2004)) | Wine-Lee, L., et al., (2004). |
| Zic2 | 1.09 | Dorsal neural tube | BMP6/7 (Lee, K. J., & Jessell, T. M. (1999)) | Ybot-Gonzalez, P., et al., (2007).<br>Sanchez-Ferras, O., et al., (2014). |
| Gadd45a | 1.08 | Mostly dorsal neural tube<br>Residual expression in ventral neural tube | BMP2 (Ijiri, K., et al. (2005)) | Kaufmann, L. T., et al., (2011). |
| Id1 | 1.04 | Roof plate and dorsal neural ectoderm<br>+ small ventral domain | BMP (Wine-Lee, L., et al., (2004)) | Wine-Lee, L., et al., (2004). |
| Cdh11 | 0.98 | Neural crest + dorsal neural tube | BMP7 (Awazu, M., et al., (2017)) | Tondeleir, D., et al., (2014).<br>Kashef, J., et al., (2009). |
| Lhx1 | 0.97 | Dorsal interneurons | Not affected by BMP4/7 (Le Dréau, G., et al., (2012)) | Le Dréau, G., et al., (2012). |
| Fgf15 | 0.95 | Dorsal regions of the di-, mes and metencephalon | - | McWhirter, J. R., et al., (1997).<br>Fischer, T., et al., (2011). |

**Supplementary Table 5.**
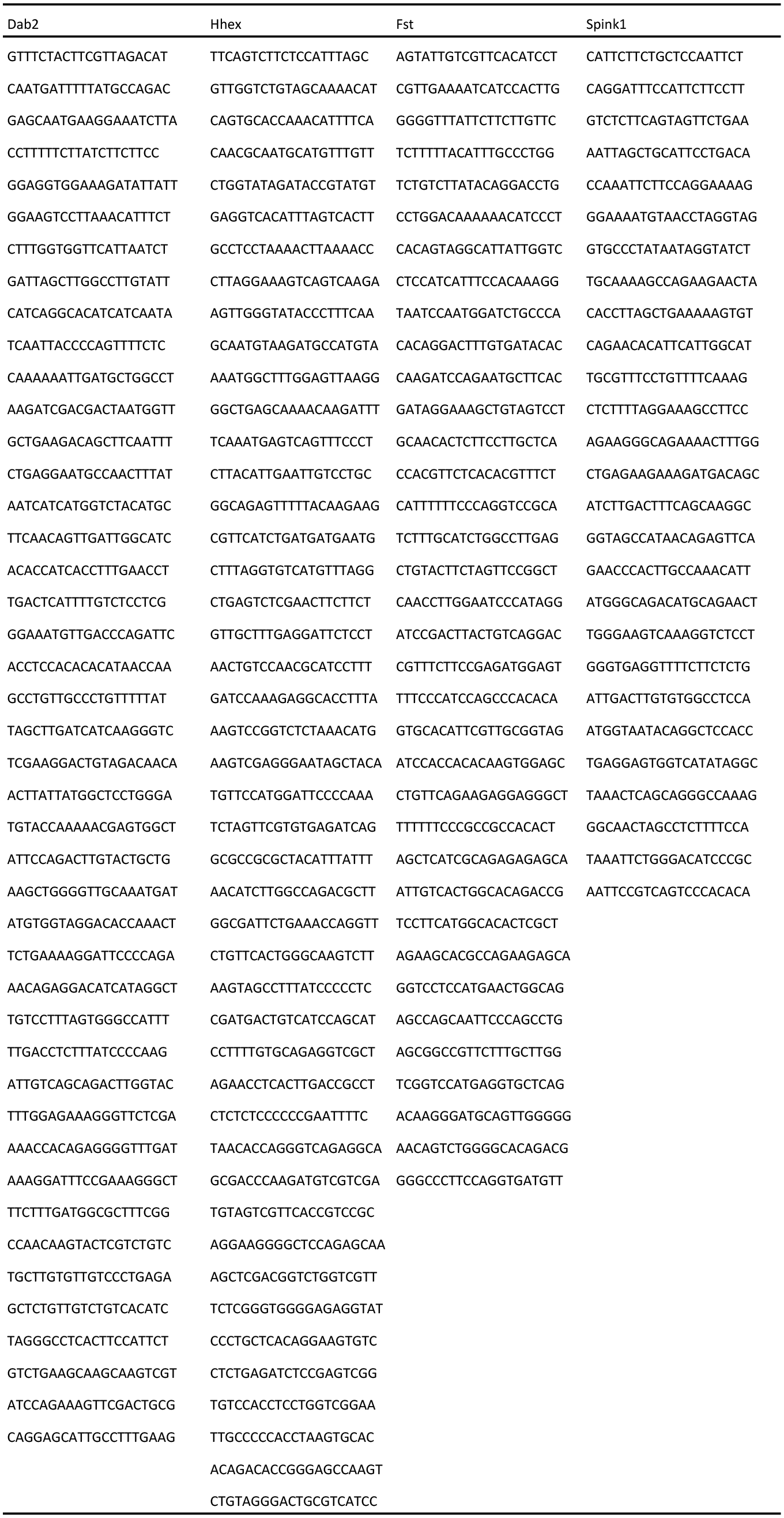
Single-molecule FISH probes.

